# Does hippocampal volume explain performance differences on hippocampal-dependent tasks?

**DOI:** 10.1101/2020.04.29.067611

**Authors:** Ian A. Clark, Anna M. Monk, Victoria Hotchin, Gloria Pizzamiglio, Alice Liefgreen, Martina F. Callaghan, Eleanor A. Maguire

## Abstract

Marked disparities exist across healthy individuals in their ability to imagine scenes, recall autobiographical memories, think about the future and navigate in the world. The importance of the hippocampus in supporting these critical cognitive functions has prompted the question of whether differences in hippocampal grey matter volume could be one source of performance variability. Evidence to date has been somewhat mixed. In this study we sought to mitigate issues that commonly affect these types of studies. Data were collected from a large sample of 217 young, healthy adult participants, including whole brain structural MRI data (0.8mm isotropic voxels) and widely-varying performance on scene imagination, autobiographical memory, future thinking and navigation tasks. We found little evidence that hippocampal grey matter volume was related to task performance in this healthy sample. This was the case using different analysis methods (voxel-based morphometry, partial correlations), when whole brain or hippocampal regions of interest were examined, when comparing different sub-groups (divided by gender, task performance, self-reported ability), and when using latent variables derived from across the cognitive tasks. Hippocampal grey matter volume may not, therefore, significantly influence performance on tasks known to require the hippocampus in healthy people. Perhaps only in extreme situations, as in the case of licensed London taxi drivers, are measurable ability-related hippocampus volume changes consistently exhibited.

**Highlights:** Evidence is mixed about whether hippocampal volume affects cognitive task performance

This is particularly the case concerning individual differences in healthy people

We collected structural MRI data from 217 healthy people

They also had widely-varying performance on cognitive tasks linked to the hippocampus

In-depth analyses showed little evidence hippocampal volume affected task performance

## 1. INTRODUCTION

People vary substantially in their ability to perform tasks related to critical aspects of cognition that enable the smooth functioning of our everyday lives. These include imagining scenes (the process of forming and visualising scene imagery in the absence of visual input), autobiographical memory (the recall of past events from one’s life), future thinking (imagining future experiences) and spatial navigation (the process of ascertaining one’s position in the environment, and planning and following a route). For example, some individuals can recollect decades-old autobiographical memories with great clarity compared to others who struggle to recall what they did last weekend (e.g. Palombo, Sheldon, & Levine, 2018). Spatial navigation can be undertaken with ease by some people while others consistently get lost (e.g. Newcombe, 2018).

Neuroimaging studies of healthy individuals performing tasks related to these cognitive functions reliably show engagement of the hippocampus (Addis, Wong, & Schacter, 2007; Barry, Barnes, Clark, & Maguire, 2018; Brown, Ross, Keller, Hasselmo, & Stern, 2010; Buckner & Carroll, 2007; Dalton, Zeidman, McCormick, & Maguire, 2018; Spiers & Maguire, 2006; Svoboda, McKinnon, & Levine, 2006). Moreover, damage to the hippocampus leads to impaired performance on similar tasks (e.g. Andelman, Hoofien, Goldberg, Aizenstein, & Neufeld, 2010; Hassabis, Kumaran, Vann, & Maguire, 2007; Maguire, Nannery, & Spiers, 2006; Rosenbaum et al., 2005; Rosenbaum et al., 2000; Scoville & Milner, 1957; Winocur & Moscovitch, 2011). The importance of the hippocampus in supporting scene imagination, autobiographical memory, future thinking and navigation, has prompted the question of whether differences in hippocampal structure could be one source of variable task performance among healthy people. While there have been numerous studies in the last two decades that have addressed this issue, typically in relation to hippocampal volume, evidence has been somewhat mixed.

Considering first scene imagination, as far as we are aware, no studies involving healthy people have yet investigated the association between hippocampal volume and the ability to construct scene imagery. Recent work documented a relationship between hippocampal grey matter volume and scene imagination performance in patients with the behavioural variant of frontotemporal dementia, a link that was not apparent in patients with Alzheimer’s disease (Wilson et al., 2020). However, in that study no direct comparison was made between the two patient groups, or between the patients and healthy controls, precluding interpretations about the association between hippocampal volume and scene imagination ability.

A larger number of studies have investigated the relationship between hippocampal volume and memory ability in healthy individuals, but with mixed results. On the one hand, “recollective” memory ability (the detailed re-experiencing of individual episodes characterized by retrieval of items in their context) has been positively associated with posterior hippocampal grey matter volume (Poppenk & Moscovitch, 2011). Moreover, a single individual identified with “Highly Superior Autobiographical Memory” was reported to have greater posterior hippocampal volume compared to matched controls (Mazzoni et al., 2019). On the other hand, group studies of individuals with superior memory capabilities, including those with Highly Superior Autobiographical Memory or those who take part in the World Memory Championships, observed no differences in hippocampal grey matter volume compared to matched controls (LePort et al., 2012; Maguire, Valentine, Wilding, & Kapur, 2003). Similarly, a meta-analysis of 33 studies found no relationship between hippocampal grey matter volume and performance on laboratory-based memory tasks (Van Petten, 2004). Whether a link exists between hippocampal volume and more naturalistic autobiographical memory ability is, therefore, unclear.

There has been limited work investigating the relationship between future thinking ability and hippocampal volume. A recent study claimed to have observed a positive relationship between participant ratings of sensory perceptual qualities in future thinking (e.g. vividness, the amount of visual details) and hippocampal grey matter volume (Yang, Chen, Zhang, Xu, & Feng, 2020). However, the peak voxel coordinates (MNI space; 39, −7.5, −10.5) and cluster were located outside of the hippocampus (Figure 1 of Yang et al., 2020). A more precise investigation into the relationship between hippocampal volume and future thinking ability is, therefore, required.

**Fig. 1.**
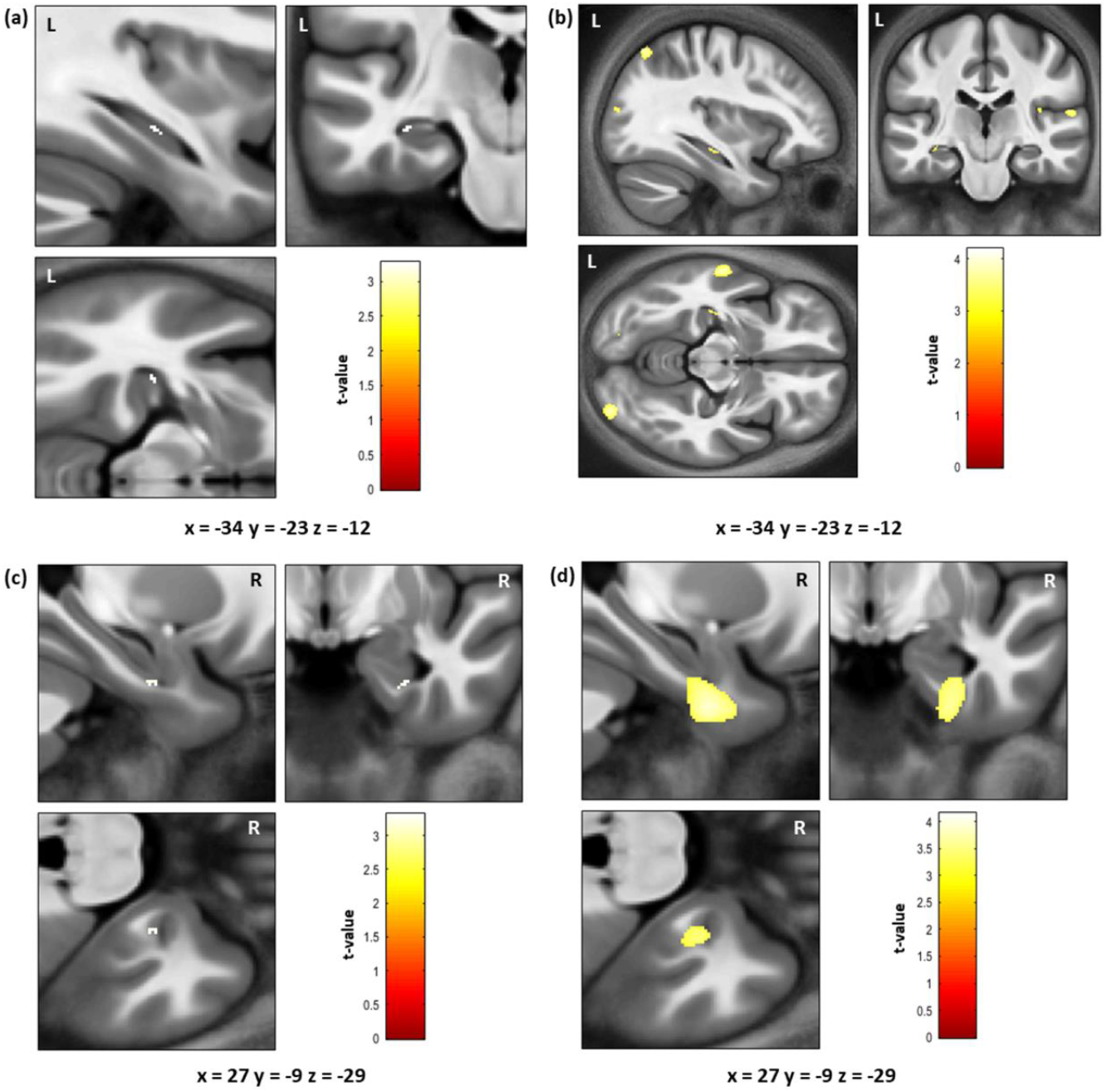
Significant results from the VBM analyses. For all panels the results are shown on the averaged brain of the whole sample (n = 217) and the coloured bar indicates the t value associated with each voxel. (a) Negative association between the Autobiographical Interview (AI) internal place sub-measure and grey matter volume limited to the bilateral hippocampus mask - 10 hippocampal voxels were identified, significant (p < 0.05) only when correcting for the posterior hippocampal mask. (b) Negative association between the AI internal place sub-measure and grey matter volume at p < 0.001 uncorrected for the whole brain. (c) Positive association between navigation route knowledge and grey matter volume limited to the bilateral hippocampus mask; 17 hippocampal voxels were identified, significant (p < 0.05) only when correcting for the anterior hippocampal mask. (d) Positive association between navigation route knowledge and grey matter volume at p < 0.001 uncorrected for the whole brain, showing that the hippocampal voxels are in fact located on the edge of a cluster centred on the right anterior fusiform and parahippocampal cortices.

A definitive link has been identified between hippocampal grey matter volume and extreme spatial navigation ability. Licensed London (UK) taxi drivers must memorise the extensive and complex layout of ~25,000 London streets and thousands of landmarks, known colloquially as acquiring “The Knowledge”. In several studies they were found to have greater posterior, and decreased anterior, hippocampal grey matter volume compared to healthy controls (Maguire et al., 2000), London bus drivers, who spend an equivalent amount of time driving but on regular routes instead of requiring a complete knowledge of London’s layout (Maguire, Woollett, & Spiers, 2006), and medical doctors, who have high levels of expertise but not primarily in the spatial domain (Woollett, Glensman, & Maguire, 2008). Moreover, longitudinal data collected before and after attempts to acquire The Knowledge identified posterior hippocampal grey matter volume enlargement within subjects, but only in those individuals who went on to qualify as London taxi drivers (Woollett & Maguire, 2011). Greater posterior, but less anterior, hippocampal grey matter volume is, therefore, reliably associated with extreme spatial navigation expertise.

In the general population, however, the relationship between hippocampal grey matter volume and navigation ability is less clear. One study involving navigation in a virtual environment found no association between hippocampal volume and navigation performance in healthy people (Maguire, Spiers, et al., 2003), a finding that has since been replicated in a larger sample of 90 individuals (Weisberg, Newcombe, & Chatterjee, 2019). However, a re-analysis of the Weisberg et al. (2019) data purports to have identified a relationship between right posterior hippocampal volume and navigation performance, but one that is dependent on self-reported navigation ability or being grouped according to navigation performance ability (He & Brown, 2020). These interactions suggested that in low performing participants a negative relationship existed between right posterior hippocampal volume and task performance. However, interactions were only observed for one of five performance measures and no multiple comparison corrections were made to the statistical threshold used to determine significance. Given the reasonably large sample size and the number of tests performed on the data (by both Weisberg et al., 2019 and He & Brown, 2020) the results of this re-analysis require replication before firm conclusions can be drawn.

Posterior hippocampal grey matter volume has been positively associated with an individual’s ability to learn new environments in the real world (Schinazi, Nardi, Newcombe, Shipley, & Epstein, 2013), in virtual reality (Brown, Whiteman, Aselcioglu, & Stern, 2014), and with first person perspective taking ability during navigation (Sherrill, Chrastil, Aselcioglu, Hasselmo, & Stern, 2018). In addition, greater posterior compared to anterior hippocampal grey matter volume has been linked to the use of “map-based” strategies, and consequently with better navigation performance (Brunec et al., 2019). By contrast, greater anterior hippocampal grey matter volume has been related to better topographical memory ability (Hartley & Harlow, 2012), path integration (Chrastil, Sherrill, Aselcioglu, Hasselmo, & Stern, 2017), and performance on a virtual reality eight arm radial maze (Bohbot, Lerch, Thorndycraft, Iaria, & Zijdenbos, 2007). Therefore, the literature on associations between navigational ability and hippocampal grey matter volume is disparate, and lacks consistency.

Overall, across scene imagination, autobiographical memory, future thinking and navigation, the relationship between hippocampal grey matter volume and task performance in healthy individuals is ambiguous. Why might this be the case?

First, the sample sizes used have typically been small, with most studies including 20-30 participants. Recent empirical work has highlighted the non-replicability of brain structure-function associations when using small samples (Kharabian Masouleh, Eickhoff, Hoffstaedter, Genon, & Alzheimer’s Disease Neuroimaging Initiative, 2019). Notably, in the navigation study where a larger sample of 90 participants was utilised, no relationship between hippocampal volume and task performance was identified (Weisberg et al., 2019). The mixed results may, therefore, simply be due to using insufficiently powered samples.

Second, there is a large variability across studies in how analyses were performed. For example, while several analyses included covariates to account for age, gender and total intracranial volume, others included covariates for only some of these variables, and numerous analyses involved no covariates at all. Given that these variables are known to be potential confounds (e.g. Ridgway et al., 2008), their inclusion (or not) as covariates could significantly influence the observed results.

Third, studies differ in terms of the level of statistical correction that was applied. For instance, some studies use more lenient statistical thresholds compared to others, and while correction for multiple comparisons is evident when testing for multiple relationships in several studies, in others it is not. Differences in statistical thresholds could not only contribute to contradictory results across the literature, but may also have increased the number of false positive findings.

Fourth, there are differences in how hippocampal anatomy was examined. Many studies investigate the hippocampus as a whole, while others divide the hippocampus into anterior and posterior portions, or focus on the volumetric ratio between the anterior and posterior segments. Different functions have been ascribed to anterior and posterior portions (Poppenk, Evensmoen, Moscovitch, & Nadel, 2013; Strange, Witter, Lein, & Moser, 2014; Zeidman & Maguire, 2016), and this could also have influenced extant results.

Finally, in some studies, differing hippocampal volume-task relationships have been identified for sub-groups of participants that were not apparent across the whole sample. However, these participant sub-groups are not always directly compared to each other making it difficult to have confidence in any reported differences. In addition, sub-group analyses are often deemed “exploratory” and “requiring replication”, but it is rare that such follow-up occurs.

In the current study we sought to overcome these issues in a comprehensive investigation of the relationship between hippocampal grey matter volume and scene imagination, autobiographical memory, future thinking and spatial navigation task performance in a healthy sample. To ensure an adequate sample size, data were collected from 217 individuals with a wide range of ability. All participants performed tasks relating to each of the aforementioned cognitive functions and hippocampal grey matter volume was derived from structural MRI scans with an isotropic voxel resolution of 0.8mm. Analyses were first performed on each task’s main outcome measure. As concerns could be raised that these four measures are over-general, for completeness, we also examined the sub-measures of each task (an additional 21 performance variables). This allowed us to investigate potential nuances that may exist in hippocampal volume-task performance relationships. As reviewed above, the mixed results in the literature made the formulation of clear hypotheses difficult. Consequently, our aim in the data analyses was to reflect the range of approaches previously pursued in the literature, and to perform a broad and inclusive evaluation of the relationship between hippocampal grey matter volume and task performance.

We have reported the main outcome measures of the four cognitive tasks in two previous papers that asked different research questions to those under consideration here. Clark et al. (2019) assessed the relationships between scene imagination, autobiographical memory, future thinking and navigation in terms of task performance. Using mediation analyses, they found that scores on the scene construction task explained the relationships between autobiographical memory, imagining the future and spatial navigation task performance. Clark & Maguire (2020) investigated how subjective questionnaire data were related to task performance. This revealed that imagination and navigation questionnaires reflected performance on their related tasks. Memory questionnaires, on the other hand, were associated with autobiographical memory vividness. In the current study, we examined whether performance on the cognitive tasks, including the main outcome measures but also their sub-measures, were related to hippocampal grey matter volume. The structural MRI data have not been published previously.

## 2. MATERIALS AND METHODS

### 2.1 Participants

Two hundred and seventeen healthy individuals were recruited, including 109 females and 108 males. The age range was restricted to 20-41 years old to limit the possible effects of ageing (mean age = 29.0 years, *SD* = 5.60). Participants had English as their first language and reported no history of psychological, psychiatric or neurological conditions. Our aim was to assess people from the general population who would not be classed as having extreme expertise in the domains of interest. Consequently, people with vocations such as taxi driving (or those training to be taxi drivers), ship navigators, aeroplane pilots, or those with regular hobbies including orienteering, or taking part in memory sports and competitions, were excluded. Participants were reimbursed £10 per hour for taking part which was paid at study completion. All participants gave written informed consent and the study was approved by the University College London Research Ethics Committee (project ID: 6743/001) and conducted in accordance with the guidelines of the Declaration of Helsinki.

### 2.2 Procedure

Participants completed the study over multiple visits. Structural MRI scans were acquired during the first visit. Cognitive testing was conducted during visits two and three. The scene imagination, autobiographical memory and future thinking tasks were completed during one visit, while the navigation tasks were performed on a separate visit. All participants completed all parts of the study.

### 2.3 Cognitive tasks

All tasks are published and were performed and scored as per their published use. Here, for the reader’s convenience, we describe each task briefly.

#### 2.3.1 Scene imagination: the scene construction task

The scene construction task (Hassabis, Kumaran, Vann, et al., 2007) measures a participant’s ability to mentally construct an atemporal visual scene, meaning that the scene is not grounded in the past or the future. Participants construct different scenes of commonplace settings. For each of seven scenes, a short cue is provided (e.g. imagine lying on a beach in a beautiful tropical bay) and the participant is asked to imagine the scene that is evoked and then describe it out loud in as much detail as possible. Recordings are transcribed for later scoring. Participants are explicitly told not to describe a memory, but to create a new atemporal scene that they have never experienced before.

The main outcome measure is the “experiential index” which is calculated for each scene and then averaged. In brief, it is composed of four elements: the content, participant ratings of their sense of presence (how much they felt like they were really there) and perceived vividness, participant ratings of the spatial coherence of the scene, and an experimenter rating of the overall quality of the scene.

For the scene construction sub-measures, we separately investigated the four categories that make up the content score, and also the spatial coherence rating.

To score the content, four categories of statement are identified; spatial references, entity presence, sensory description and thoughts/emotions/actions. The spatial reference category encompasses statements regarding the relative position of entities within the environment or directions relative to the participant’s vantage point. The entity category is a count of how many distinct entities (e.g., objects, people, animals) were mentioned. The sensory descriptions category consists of any statements describing (in any modality) properties of an entity or the environment in general. Finally, the thoughts/emotions/actions category covers any introspective thoughts or emotional feelings as well as the thoughts, intentions, and actions of other entities in the scene. The final score of each category is the average across the seven scenes.

The imagined scenes are also examined using the spatial coherence index. After each scene is mentally constructed, participants are presented with a set of 12 statements, each providing a possible qualitative description of the imagined scene. Participants are instructed to indicate which statements they felt accurately described their construction, identifying as many or as few as they thought appropriate. Eight of the statements indicate that aspects of the scene were integrated (e.g. “I could see the whole scene in my mind’s eye”), whereas four indicate that aspects of the scene were fragmented (e.g. “It was a collection of separate images”). One point is awarded for each integrated statement selected and one point taken away for each fragmented statement. This yields a score between –4 and +8 that is then normalised around zero. Any construction with a negative score is considered to be incoherent and fragmented and scored at 0 so as not to over-penalise fragmented descriptions. The final spatial coherence index was the average score of the seven scenes, ranging between 0 (totally fragmented) and +6 (completely integrated).

Double scoring was performed on 20% of the data. We took the most stringent approach to identifying across-experimenter agreement. Inter-class correlation coefficients, with a two-way random effects model looking for absolute agreement indicated excellent agreement among the experimenters (minimum score of 0.9; see Supplementary Methods Table S1). For reference, a score of 0.8 or above is considered excellent agreement beyond chance.

#### 2.3.2 Autobiographical memory: the autobiographical interview

In the autobiographical interview (AI; Levine, Svoboda, Hay, Winocur, & Moscovitch, 2002) participants are asked to provide autobiographical memories from a specific time and place over four time periods – early childhood (up to age 11), teenage years (aged from 11-17), adulthood (from age 18 years to 12 months prior to the interview; two memories are requested) and the last year (a memory from the last 12 months). Recordings are transcribed for later scoring.

The AI has two main outcome measures; the number of “internal” and “external” details included in the description of an event. Importantly, these two scores represent different aspects of autobiographical memory recall. Internal details are those describing the event in question (i.e. episodic details), and were of primary interest here. External details describe semantic information concerning the event, or non-event information. Internal events are, therefore, thought to be hippocampal-dependent, while external events are not. The two AI scores are obtained by separately averaging performance for the internal and external details across the five autobiographical memories.

For the AI sub-measures, we examined the five separate categories that comprise the internal details outcome measure, as well as considering AI vividness ratings. While external details can also be split into component categories, too few details were provided by the participants to assess each of these individually.

Internal details are composed of event, place, time, perceptual, and thoughts/emotions categories. The event category contains details regarding happenings or the unfolding of the story, including individuals present, actions and reactions. The time category refers to any details of the year, season, day or time of day wherein the event occurred. The place category contains details that localise the event, both at the general level (e.g. a city), and more specifically (e.g. to parts of a room). The perceptual category involves descriptions (in any modality) of any aspect of the event. Finally, the thoughts/emotions category includes descriptions of emotional states, thoughts or implications. The final score of each category is the average across the five memories.

AI vividness ratings were examined given a recent finding that responses on various memory questionnaires were associated with AI vividness, suggesting that vividness may be a key process in autobiographical memory recall (Clark & Maguire, 2020). Vividness ratings are collected for each memory in response to the question “How clearly can you visualize this event?” on a 6-point scale from 1 (vague memory, no recollection) to 6 (extremely clear as if it’s happening now). An overall vividness rating was the average of the vividness ratings provided for each autobiographical memory.

Double scoring was performed on 20% of the data, and there was excellent agreement across the experimenters (minimum score of 0.81; see Supplementary Methods Table S2).

#### 2.3.3 Future thinking

The future thinking task (Hassabis, Kumaran, Vann, et al., 2007) follows the same procedure as the scene construction task, but requires participants to imagine three plausible future scenes involving themselves (an event at the weekend; next Christmas; the next time they will meet a friend). There are two main differences between the future thinking task and the scene construction task. First, unlike scenes in the scene construction task, scenes in the future thinking task involve ‘mental time travel’ to the future, so they have a clear temporal dimension. Second, the cues for the scene construction task are somewhat more specific than those employed in the future thinking task (see Hurley, Maguire, & Vargha-Khadem, 2011 for more on this issue).

The scoring procedures for both the main outcome measure and the sub-measures are the same as for the scene construction task. Double scoring identified excellent agreement across the experimenters (minimum score of 0.88; see Supplementary Methods Table S3).

#### 2.3.4 Navigation

Navigation ability was assessed using the paradigm described by (Woollett & Maguire, 2010). A participant watches movie clips of two overlapping routes through an unfamiliar real town (Blackrock, Dublin, Ireland) four times.

The main outcome measure for navigation performance is calculated by combining the scores from the five tasks used to assess navigational ability. The sub-measures are the performance scores of each individual task.

The tasks were as follows. First, was a movie clip recognition task; following each viewing of the route movies, participants are shown four short movie clips – two from the actual routes, and two distractors. Participants indicate whether they have seen a movie clip before or not. Second, after all four route viewings are completed, recognition memory for scenes from the routes is tested. The third task, proximity judgements, involves assessing knowledge of the spatial relationships between landmarks from the routes. Fourth, the route knowledge task, requires participants to place scene photographs from the routes in the correct order as if travelling through the town. Finally, the sketch map task involves participants drawing a sketch map of the two routes including as many landmarks as they can remember. Sketch maps are scored in terms of the number of road segments, road junctions, correct landmarks, landmark positions, the orientation of the routes and an overall map quality score from the experimenters. Double scoring was performed on 20% of the sketch maps, and there was excellent agreement among the experimenters (minimum of 0.89; see Supplementary Methods Table S4).

### 2.4 Self-report measures

#### 2.4.1 Plymouth Sensory Imagery Questionnaire, Appearance subscale

(Andrade, May, Deeprose, Baugh, & Ganis, 2014). The PSIQ Appearance subscale assesses the vividness of visual imagery. The subscale requires participants to imagine three scenarios: a bonfire, a sunset, and a cat climbing a tree. They then rate the visual image they generated on an 11-point scale from 0 (no image at all) to 10 (vivid as real life). Scores on the three scenarios are summed to create a total score out of 30. Responses on the PSIQ have previously been associated with performance on the scene construction, autobiographical memory, and future thinking tasks (Clark & Maguire, 2020).

#### 2.4.2 Santa Barbara Sense of Direction Scale

(Hegarty, Richardson, Montello, Lovelace, & Subbiah, 2002). The SBSODS assesses spatial and navigational abilities, preferences and experiences. Fifteen statements are presented, with participants indicating their level of agreement with each statement. Ratings are made on a 7-point scale from 1 (strongly agree) to 7 (strongly disagree). Seven statements are positively coded (“I am very good at giving directions”) and eight are reverse scored (“I don’t have a very good “mental map” of my environment”). Scores are summed across the 15 statements to create a total score out of 105 (where a low score reflects good navigation ability). Responses on the SBSODS have been associated with multiple measures of navigation, including the spatial navigation task used in the current study (Clark & Maguire, 2020; Hegarty et al., 2002).

### 2.5 Statistical analyses of the behavioural data

Data were summarised using means and standard deviations, calculated in SPSS v22. There were no missing data, and no data needed to be removed from any analysis.

### 2.6 MRI data acquisition

Three MRI scanners were used to collect the structural neuroimaging data. All scanners were Siemens Magnetom TIM Trio systems with 32 channel head coils and were located at the same imaging centre, running the same software. The sequences were loaded identically onto the individual scanners. Participant set-up and positioning followed the same protocol for each scanner.

Whole brain volumetric images had an isotropic resolution of 800μm × 800μm × 800μm and were obtained using magnetisation transfer (MT) saturation maps derived from a multi-parameter mapping (MPM) quantitative imaging protocol (Callaghan et al., 2015; Weiskopf et al., 2013). This protocol consisted of the acquisition of three multi-echo gradient acquisitions with either proton density (PD), T1 or MT weighting. Each acquisition had a repetition time, TR, of 25 ms. PD weighting was achieved with an excitation flip angle of 6^0^, which was increased to 21^0^ to achieve T1 weighting. MT weighting was achieved through the application of a Gaussian RF pulse 2 kHz off resonance with 4ms duration and a nominal flip angle of 220^0^. This acquisition had an excitation flip angle of 6^0^. The field of view was 256mm head-foot, 224mm anterior-posterior (AP), and 179mm right-left (RL). The multiple gradient echoes per contrast were acquired with alternating readout gradient polarity at eight equidistant echo times ranging from 2.34 to 18.44ms in steps of 2.30ms using a readout bandwidth of 488 Hz/pixel. Only six echoes were acquired for the MT weighted volume to facilitate the off-resonance pre-saturation pulse within the TR. To accelerate the data acquisition, partially parallel imaging using the GRAPPA algorithm was employed in each phase-encoded direction (AP and RL) with forty reference lines and a speed up factor of two. Calibration data were also acquired at the outset of each session to correct for inhomogeneities in the RF transmit field (Lutti, Hutton, Finsterbusch, Helms, & Weiskopf, 2010; Lutti et al., 2012).

### 2.7 MRI data pre-processing

The data from the MPM protocol were processed for each participant using the hMRI toolbox (Tabelow et al., 2019) within SPM12 (www.fil.ion.ucl.ac.uk/spm). The default toolbox configuration settings were used, with the exception that correction for imperfect spoiling was additionally enabled. The output MT saturation map used for the volumetric analyses quantified the degree of saturation of the steady state signal having accounted for spatially varying T_1_ times and RF field inhomogeneity (Helms, Dathe, & Dechent, 2008; Weiskopf et al., 2013).

Each participant’s MT saturation map was then segmented into grey matter probability maps using the unified segmentation approach (Ashburner & Friston, 2005), but with no bias field correction (since the MT saturation map does not suffer from any bias field modulation). Inter-subject registration was performed using DARTEL, a nonlinear diffeomorphic algorithm (Ashburner, 2007), as implemented in SPM12. This algorithm estimates the deformations that best align the tissue probability maps by iteratively registering them with their average. The resulting DARTEL template and deformations were used to normalize the tissue probability maps to the stereotactic space defined by the Montreal Neurological Institute (MNI) template (at 1 x 1 x 1mm resolution), with an isotropic Gaussian smoothing kernel of 8mm full width at half maximum (FWHM).

### 2.8 Primary VBM analyses

Our primary analyses investigating the associations between cognitive task performance and hippocampal grey matter volume were performed using voxel-based morphometry (VBM) (Ashburner & Friston, 2000; Mechelli, Price, Friston, & Ashburner, 2005).

First, we investigated the relationship between hippocampal volume and the main outcome measures for each of the cognitive tasks assessing scene imagination, autobiographical memory, future thinking and navigation. We then examined the associations between hippocampal volume and each of the sub-measures from these tasks.

For each performance measure, statistical analyses were carried out using multiple linear regression models. Six regressors were included in each model, the cognitive task performance measure of interest and five covariates: one each for age, gender and total intracranial volume, and two regressors to model the effects of the different scanners (only two regressors are required because one scanner is taken as the baseline, with the two regressors then modelling any effects that are different between the other scanners and the baseline scanner). The dependent variable was the smoothed and normalised grey matter volume.

#### 2.8.1 Whole brain VBM

Analyses were first carried out voxel-wise across whole brain grey matter using an explicitly defined mask which was generated by averaging the smoothed grey matter probability maps in MNI space across all subjects. Voxels for which the grey matter probability was below 80% were excluded from the analysis. Two-tailed t-tests were used to investigate the relationships between cognitive task performance and grey matter volume, with statistical thresholds applied at p < 0.05 family-wise error (FWE) corrected for the whole brain, and a minimum cluster size of 5 voxels.

As our main focus was on the relationship between cognitive task performance and hippocampal grey matter volume, in the main text we report only findings pertaining to the hippocampus. However, performing analyses at the whole brain level meant that we could apply the recommended statistical threshold to the analyses (Nichols et al., 2017; Poldrack et al., 2008). Consequently any significant relationships identified would be supported by the strongest evidence. In addition, performing analyses at the whole brain level also allowed us to investigate whether any non-hippocampal brain regions were associated with cognitive task performance – these results are reported in the Supplementary Results, although there were very few.

#### 2.8.2 Hippocampal ROI VBM

Following the whole brain analysis, we focused on the hippocampus using anatomical hippocampal masks. This allowed us to investigate whether any weaker relationships existed between hippocampal grey matter volume and task performance that did not reach the statistical threshold required for the whole brain analyses. The masks were manually delineated on the group-averaged MT saturation map in MNI space (1mm × 1mm × 1 mm) using ITK-SNAP (www.itksnap.org). As in Poppenk & Moscovitch (2011) and Brunec et al. (2019), the anterior hippocampus was delineated using an anatomical mask that was defined in the coronal plane and proceeded from the first slice where the hippocampus can be observed in its most anterior extent until the final slice of the uncus. The posterior hippocampus was defined from the first slice following the uncus until the final slice of observation in its most posterior extent (see Dalton, Zeidman, Barry, Williams, & Maguire, 2017 for more details). The whole hippocampal mask comprised the combination of the anterior and posterior masks. The masks were examined separately for the left and right hippocampus, and as bilateral masks.

Two-tailed t-tests were used to investigate the relationships between cognitive task performance and grey matter volume within the hippocampal masks. Voxels were regarded as significant when falling below an initial whole brain uncorrected voxel threshold of p < 0.001, and then a small volume correction threshold of p < 0.05 FWE corrected for each mask, with a minimum cluster size of 5 voxels. Given that all significant results observed using the anterior and posterior unilateral masks were also evident using the bilateral masks, we report the results of the bilateral masks. Where significant results within the masks were identified, the whole brain analysis was re-examined with the threshold set to p < 0.001 uncorrected to assess whether the effects identified within the hippocampal masks were due to leakage from adjacent brain regions.

### 2.9 Auxiliary analyses using extracted hippocampal volumes

Following our main analyses, we performed a series of auxiliary investigations to further scrutinise potential relationships between hippocampal grey matter volume and scene imagination, autobiographical memory, future thinking and navigation task performance.

The basis of these auxiliary analyses was the hippocampal grey matter volume for each participant that was extracted using ‘spm_summarise’. The whole, anterior and posterior anatomical hippocampal masks (described above) were applied to each participant’s smoothed and normalised grey matter volume maps, and the total volume within each mask was extracted. It has been suggested that the posterior:anterior hippocampal volume ratio shows stronger relationships to memory and navigation performance than raw anterior and posterior hippocampal volumes alone (Brunec et al., 2019; Poppenk & Moscovitch, 2011). Therefore, in line with these studies, we also calculated each participant’s posterior:anterior hippocampal volumetric ratio. We did this by dividing the posterior hippocampal volume by the anterior hippocampal volume, providing a ratio whereby values greater than 1 were indicative of a larger posterior over anterior hippocampus and those less than 1 indicated a larger anterior over posterior hippocampus.

Overall, across all our auxiliary investigations (described in detail below) we examined the relationships between four different measures of hippocampal grey matter volume (whole, anterior, posterior, ratio) for each of our 26 task performance measures. Given the extensive nature of these analyses, we needed to ensure that appropriate correction was made to the statistical thresholds. Typically, FWE corrections (such as Bonferroni) are employed in this regard. However, FWE correction acts to ensure that the probability of a *single* false positive result remains at the critical alpha (typically p < 0.05); a highly effective method if the need to avoid false positives is high. However, this level of control comes at a price, especially when controlling for a large number of statistical tests, as it means that the effects of any single variable have to be particularly large to reach the adjusted significance level. Thus, while false positives are avoided, the risk of incorrectly rejecting a true result is introduced.

An alternative method of statistical correction is to control the false discovery rate (Benjamini & Hochberg, 1995) which influences the *proportion* of significant results identified that are false positives. Therefore, a FDR of p < 0.05 allows for 5% false positive results across the tests performed. Controlling for the FDR is, therefore, more lenient than performing FWE corrections, as it accommodates the potential for a greater number of false positive results but, by doing so, also reduces the chances of incorrectly rejecting a true result. Applying our statistical correction in terms of the FDR provided, therefore, a compromise between the need to adjust our statistical thresholds for multiple comparisons, while not removing all possibility of identifying true relationships among the data features.

Across the auxiliary analyses, the FDR was set to p < 0.05 for each family of tests. A family was defined as all the tests performed for each set of auxiliary analyses for one cognitive task. For example, when testing for relationships between hippocampal volume and scene construction task performance, we had 4 measures of hippocampal volume (whole, anterior, posterior, ratio) and 6 measures of performance (experiential index, spatial references, entities present, sensory descriptions, thoughts/emotions/actions, spatial coherence), totalling 24 separate statistical tests. The FDR was thus set to p < 0.05 for these 24 tests.

The following auxiliary analyses were performed in SPSS v25 with FDR thresholding and Benjamini-Hochberg adjusted p values calculated using the resources provided by McDonald (2014).

#### 2.9.1 Partial correlations using extracted hippocampal volumes

We performed partial correlation analyses between each of the hippocampal volumetric measures (whole, anterior, posterior, ratio) and each of our measures of task performance. As with the primary VBM analyses, age, gender, total intracranial volume and MRI scanner were included as covariates.

#### 2.9.2 Sub-group analyses using extracted hippocampal volumes

##### 2.9.2.1 Effects of gender

To directly investigate the effects of gender, we divided the sample into male (n = 108) and female (n = 109) participants. We first performed separate partial correlations for the two participant groups for each of the hippocampal volumetric measures (whole, anterior, posterior, ratio) and each of our measures of task performance. Age, total intracranial volume and MRI scanner were included as covariates. If a significant volume-task relationship was identified in either the male or female group, our intention was to then compare the male and female relationships directly. However, this was never required (see section 3.3.2).

##### 2.9.2.2 Median split: direct comparison

Participants were first allocated to groups depending on their performance on a task compared to the median score for that task. If their score was less than or equal to the median value they were assigned to the low performing group, scores greater the median resulted in allocation to the high performing group. Due to the distribution of performance scores, the navigation movie clip recognition test could not be divided into groups (as most participants performed at ceiling) and so was not included in these analyses. Our four measures of hippocampal volume were then compared between the low and high performing groups for each task using univariate ANCOVAs with the hippocampal volume measure as the dependent variable, performance group as the main predictor and age, gender, total intracranial volume and MRI scanner as covariates.

##### 2.9.2.3 Median split: partial correlations

We next investigated if different associations between hippocampal volume and cognitive task scores existed in the high and low performing participants. As before, because the navigation movie clip recognition test could not be divided into groups it was not included in these analyses. For each task, partial correlations were performed separately for the two groups between the hippocampal volume measures and the task performance measures, with age, gender, total intracranial volume and MRI scanner included as covariates. If a significant relationship was identified in either the low or high performance group, our intention was to then compare the group correlations directly. However, this was never required (see section 3.3.3).

##### 2.9.2.4 Best versus worst

The best and worst performers for each task were identified. This was approximately the top and bottom 10% (n ~ 20 in each group). The exact number of participants allocated to each group varied for each task depending on the distribution of performance scores and to ensure the number of participants in each group was as similar as possible. As the navigation movie clip recognition test could not be split into approximately equal groups of best and worst performers it was not included in these analyses. Our four measures of hippocampal volume were then compared between the best and worst performing participants for each task using univariate ANCOVAs with the hippocampal volume measure as the dependent variable, performance group as the main predictor and age, gender, total intracranial volume and MRI scanner as covariates.

##### 2.9.2.5 Effects of self-reported ability

Self-reported ability was assessed using the PSIQ (Andrade et al., 2014) for scene imagination, autobiographical memory and future thinking, and the SBSODS (Hegarty et al., 2002) for navigation. Our aim here was to investigate whether the interaction between self-reported ability and hippocampal volume was associated with task performance, following the method of He & Brown (2020). As such, eight interaction terms were created: responses on the PSIQ and SBSODS by the four measures of hippocampal grey matter volume. Partial correlation analyses were then performed between the interaction terms and task performance, with age, gender, total intracranial volume and MRI scanner as covariates as in the previous analyses, with the additional covariates of the relevant questionnaire and hippocampal volume measurement to ensure that any identified relationships were due to the interaction of self-report and volume and not either individually.

### 2.10 Multivariate analyses using latent variables

The analyses detailed above are all univariate and task-based. A multivariate approach using latent variables derived from across the cognitive data may provide additional insights into associations between task performance and hippocampal grey matter volume. For example, this method offered increased statistical power and enabled data driven investigations, eschewing a reliance on conceptually driven cognitive constructs, and so had the potential to increase the generalisability of findings beyond the specific tasks in question.

To identify latent variables across the cognitive data, two Principal Component Analyses (PCA) were performed using SPSS v25 with varimax rotation and an eigenvalue cut-off of 1. The first PCA investigated possible latent variables across the four main outcome measures of the scene imagination, autobiographical memory, future thinking and navigation tasks, and the second PCA examined all 21 task sub-measures. Factor scores for the identified latent variables were extracted using the Anderson-Rubin method. The relationships between the latent variable factor scores and hippocampal grey matter volume were examined using the VBM and partial correlation approaches described above.

## 3. RESULTS

### 3.1 Cognitive task performance

A summary of the outcome measures for the cognitive tasks is shown in Table 1. A wide range of scores was obtained for all variables with the exception of navigation movie clip recognition, where performance was close to ceiling.

**Table 1.**
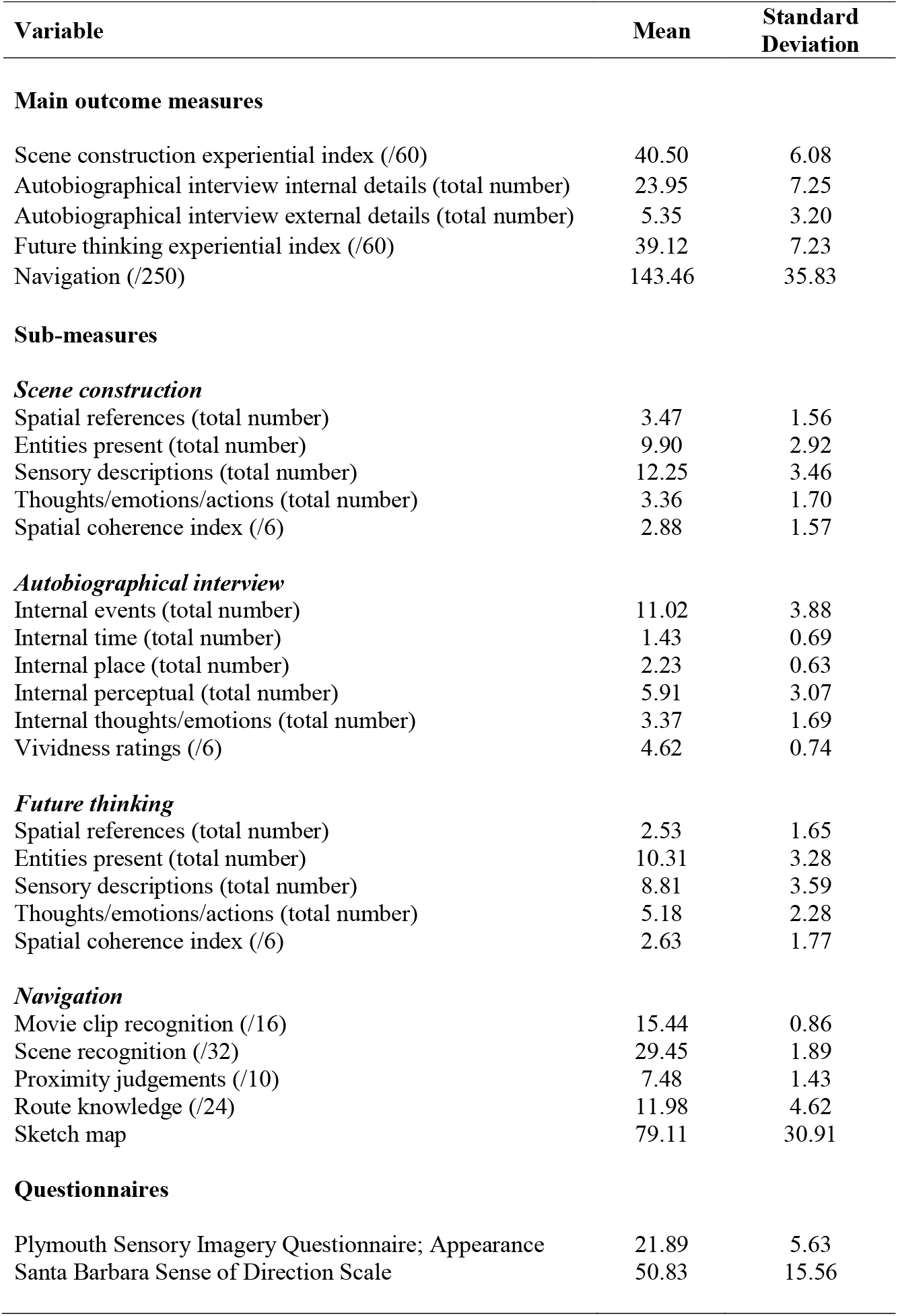
Means and standard deviations for the main outcome measures and sub-measures of the cognitive tasks and questionnaires.

| Variable | Mean | Standard Deviation |
| --- | --- | --- |
| **Main outcome measures** |  |  |
| Scene construction experiential index (/60) | 40.50 | 6.08 |
| Autobiographical interview internal details (total number) | 23.95 | 7.25 |
| Autobiographical interview external details (total number) | 5.35 | 3.20 |
| Future thinking experiential index (/60) | 39.12 | 7.23 |
| Navigation (/250) | 143.46 | 35.83 |
| **Sub-measures** |  |  |
| ***Scene construction*** |  |  |
| Spatial references (total number) | 3.47 | 1.56 |
| Entities present (total number) | 9.90 | 2.92 |
| Sensory descriptions (total number) | 12.25 | 3.46 |
| Thoughts/emotions/actions (total number) | 3.36 | 1.70 |
| Spatial coherence index (/6) | 2.88 | 1.57 |
| ***Autobiographical interview*** |  |  |
| Internal events (total number) | 11.02 | 3.88 |
| Internal time (total number) | 1.43 | 0.69 |
| Internal place (total number) | 2.23 | 0.63 |
| Internal perceptual (total number) | 5.91 | 3.07 |
| Internal thoughts/emotions (total number) | 3.37 | 1.69 |
| Vividness ratings (/6) | 4.62 | 0.74 |
| ***Future thinking*** |  |  |
| Spatial references (total number) | 2.53 | 1.65 |
| Entities present (total number) | 10.31 | 3.28 |
| Sensory descriptions (total number) | 8.81 | 3.59 |
| Thoughts/emotions/actions (total number) | 5.18 | 2.28 |
| Spatial coherence index (/6) | 2.63 | 1.77 |
| ***Navigation*** |  |  |
| Movie clip recognition (/16) | 15.44 | 0.86 |
| Scene recognition (/32) | 29.45 | 1.89 |
| Proximity judgements (/10) | 7.48 | 1.43 |
| Route knowledge (/24) | 11.98 | 4.62 |
| Sketch map | 79.11 | 30.91 |
| **Questionnaires** |  |  |
| Plymouth Sensory Imagery Questionnaire; Appearance | 21.89 | 5.63 |
| Santa Barbara Sense of Direction Scale | 50.83 | 15.56 |

### 3.2 Primary VBM analyses

#### 3.2.1 Whole brain VBM

As our main focus was on the relationship between cognitive task performance and hippocampal grey matter volume, here we report only findings pertaining to the hippocampus – any regions identified outside the hippocampus are reported in the Supplementary Results.

No significant relationships between cognitive task performance and hippocampal grey matter volume were identified for any of the main outcome measures of the tasks assessing scene imagination, autobiographical memory, future thinking or navigation. This was also the case for the sub-measures of these tasks.

#### 3.2.2 Hippocampal ROI VBM

No relationships between cognitive task performance and hippocampal grey matter volume were identified using any of the hippocampal masks for the main outcome measures from the tasks examining scene imagination, autobiographical memory, future thinking or navigation.

No associations were observed between performance on the scene construction task sub-measures and hippocampal grey matter volume using any of the hippocampal masks.

For the AI sub-measures, one potential relationship was identified. When using the posterior hippocampal mask, a cluster of 10 voxels, located towards the anterior portion of the posterior mask in the left hippocampus, was found to be negatively associated with the number of internal place references (Figure 1a; peak coordinates = −34, −23, −12, peak t = 3.28, p _FWE posterior hippocampus ROI corrected_ = 0.046). This relationship, however, was not significant when correcting for the whole hippocampus mask (p _FWE whole hippocampus ROI corrected_ = 0.069). Reducing the statistical threshold to p < 0.001 uncorrected for the whole brain confirmed the cluster was confined to the hippocampus and not due to leakage from adjacent brain regions (Figure 1b). However, visualisation at this lower threshold also demonstrated that the identified relationship was not unique to the hippocampal voxels, with a number of other brain regions also being negatively associated with the AI internal place references at this liberal statistical threshold.

For the future thinking sub-measures, no associations were observed between cognitive task performance and hippocampal grey matter volume.

Considering the navigation sub-measures, one potential relationship was identified. When using the anterior hippocampal mask, a cluster of 17 voxels, located at the edge of the mask in the right hippocampus, was found to be positively associated with performance on the navigation route knowledge sub-measure (Figure 1c; peak coordinates = 27 −8 −29, peak t = 3.31, p _FWE anterior hippocampus ROI corrected_ = 0.028). This relationship, however, was not significant when correcting for the whole hippocampus mask (p _FWE whole hippocampus ROI corrected_ = 0.062). Reducing the statistical threshold to p < 0.001 uncorrected for the whole brain clearly indicated that the identified hippocampal voxels were located on the edge of a larger cluster that was in fact centred on the right anterior fusiform and parahippocampal cortices (Figure 1d, cluster size = 1557, peak coordinates = 29 −5 −37, peak t = 3.87).

### 3.3 Auxiliary analyses using extracted hippocampal volumes

We next performed a series of auxiliary analyses to further investigate possible relationships between cognitive task performance and hippocampal grey matter volume. The latter was extracted for each participant using the predefined anatomical hippocampal masks. The mean grey matter volume for the whole hippocampus was 4456.49 mm^3^ (*SD* = 373.74 mm^3^), for the anterior mask was 2303.70 mm^3^ (*SD* = 215.07 mm^3^) and for the posterior mask was 2152.79 mm^3^ (*SD* = 182.66 mm^3^), with a mean posterior/anterior ratio of 0.94 (*SD* = 0.057), where a ratio less than 1 indicates greater anterior relative to posterior hippocampal volume.

#### 3.3.1 Partial correlations using extracted hippocampal volumes

No relationships were identified between hippocampal grey matter volume and performance for any of the main or sub-measures of the tasks assessing scene imagination, autobiographical memory, future thinking or navigation (see Supplementary Results Table S5 for details). These partial correlation findings, therefore, support those of the primary VBM analyses, as well as extending this to include the posterior:anterior hippocampal volume ratio.

#### 3.3.2 Sub-group analyses using extracted hippocampal volumes

##### 3.3.2.1 Effects of gender

We next divided the sample into male and female participants. No relationships between cognitive task performance and hippocampal grey matter volume were observed in either group for any of the main or sub-measures (see Supplementary Results Tables S6 and S7 for details). As no relationships were detected in either group, direct comparison of the relationships was not relevant.

##### 3.3.2.2 Median split: direct comparison

Details of the groups that were created when the sample was divided by median splits of performance are provided in the Supplementary Results (Tables S8). Comparing hippocampal grey matter volume measurements between the two groups determined by their median performance on each main and sub-measure found no differences between the groups for any task (see Supplementary Results Table S9 for details).

##### 3.3.2.3 Median split: partial correlations

No significant relationships between hippocampal grey matter volume and cognitive task main and sub-measures were identified in either the low or high performing groups (see Supplementary Results Tables S10 and S11 for details). As no relationships were detected in either group, direct comparison of the relationships was not relevant.

##### 3.3.2.4 Best versus worst

Details of the groups that were created when the sample was divided in terms of best and worst performers are provided in the Supplementary Results (Tables S12). Taking the best and worst performing participants for each cognitive task main and sub-measure and comparing the hippocampal grey matter volume measurements between the two groups also showed no significant differences between the groups (see Supplementary Results Table S13 details).

##### 3.3.2.5 Effects of self-reported ability

The results of these analysis are provided in the Supplementary Results (Table S14).

No significant associations were observed between any of the scene construction task main and sub-measures and the interaction between self-reported ability (determined by the PSIQ) and hippocampal grey matter volume.

For autobiographical memory, a potential relationship was identified for AI vividness, but no other main or sub-measure. For AI vividness, an association was observed between AI vividness ratings and the PSIQ by whole hippocampus volume interaction (r = 0.24, p _FDR corrected_ = 0.016), and with the PSIQ by posterior hippocampal volume interaction (r = 0.22, p _FDR corrected_ = 0.013). This suggests that self-reported imagery ability may be influencing the relationship between hippocampal volume and AI vividness ratings.

To better understand these interactions, simple slopes and region of significance analyses were performed using the interactions toolbox (Long, 2019) in R 3.6.1 (R Core Team, 2017), with regions of significance calculated using the Johnson-Neyman technique (Johnson & Fay, 1950).

First, we focused on the relationship between AI vividness and the PSIQ by whole hippocampus interaction. Repeating the partial correlation analysis as a regression (with AI vividness as the dependent variable, PSIQ, whole hippocampal volume and the PSIQ by whole hippocampal volume interaction as predictors, and age, gender, scanner and ICV as covariates), identified that for the PSIQ by whole hippocampus volume interaction, the unstandardized beta value was 0.0000710. Performing simple slopes to decompose this relationship identified that for individuals who were 1 standard deviation below the mean PSIQ score, the unstandardized beta value was −0.000859 (t = −4.15, p _FDR corrected_ < 0.001). For individuals with a mean PSIQ score, the unstandardized beta value was −0.000459 (t = −2.78, p _FDR corrected_ = 0.009), while for individuals who were 1 standard deviation above the mean PSIQ score, the unstandardized beta value was −0.0000583 (t = −0.28, p _FDR corrected_ = 0.92). As such, in individuals with low PSIQ scores, there was a negative relationship between whole hippocampal volume and AI vividness that was not evident in individuals with high PSIQ scores. A region of significance analysis found that the negative relationship existed in individuals with PSIQ scores less than 23.65.

We then investigated the relationship between AI vividness and the PSIQ by posterior hippocampus interaction. Repeating the partial correlation analysis as a regression (with AI vividness as the dependent variable, PSIQ, posterior hippocampal volume and the PSIQ by posterior hippocampal volume interaction as predictors, and age, gender, scanner and ICV as covariates), identified that for the PSIQ by posterior hippocampus volume interaction, the unstandardized beta value was 0.000157. Performing simple slopes identified that for individuals who were 1 standard deviation below the mean PSIQ score, the unstandardized beta value was −0.00180 (t = −4.53, p _FDR corrected_ < 0.001). For individuals with a mean PSIQ score, the unstandardized beta value was −0.000919 (t = −3.24, p _FDR corrected_ = 0.003), while for individuals who were 1 standard deviation above the mean PSIQ score, the unstandardized beta value was −0.0000343 (t = −0.098, p _FDR corrected_ = 0.92). As with the whole hippocampal volume interaction, in individuals with low PSIQ scores, there was a negative relationship between posterior hippocampal volume and AI vividness that was not apparent in individuals with high PSIQ scores. A region of significance analysis found that the negative association existed in individuals with PSIQ scores less than 24.13.

For all of the future thinking and navigation main and sub-measures, no significant findings were observed when investigating the association between cognitive task performance and the interaction between self-reported ability and hippocampal grey matter volume.

### 3.4 Multivariate analyses using latent variables

The detailed outcomes of the multivariate analyses can be found in the Supplementary Results. In summary, performing a PCA with the four main outcome measures of the scene imagination, autobiographical memory, future thinking and navigation tasks resulted in a single component that explained 56.05% of the variance (Supplementary Results Table S15). Conducting a VBM analysis with this latent variable identified no significant relationship between hippocampal grey matter volume and the latent variable factor scores, both at the whole brain level and when using the hippocampal ROIs. Performing partial correlations between the extracted hippocampal grey matter volumes and the latent variable factor scores also identified no significant associations (Supplementary Results).

The second PCA using all 21 task sub-measures identified 6 components that explained 65.01% of the variance (Supplementary Results Tables S16 and S17, and Figure S1). One reassuring finding of this PCA was the broad correspondence of the components to the cognitive tasks. For example, all of the navigation sub-measures loaded onto Component 2, and all of the autobiographical memory sub-measures, excluding internal time, were placed in Component 5. The scene construction and future thinking sub-measures showed greater interrelatedness, which is not surprising given the close similarity of the task requirements. Component 1 comprised the content sub-measures of both tasks, and Component 3 contained their spatial coherence indices, as well as autobiographical memory vividness ratings. Component 4 encompassed the thoughts and emotions elements of scene construction, future thinking and autobiographical memory. Finally, Component 6 corresponded predominantly to the autobiographical memory sub-measure of internal time. We then examined whether there were any relationships between these latent constructs and hippocampal grey matter volume. Using VBM (at both the whole brain level and in the anatomical ROIs) or when performing partial correlations using the extracted hippocampal grey matter volumes (Supplementary Results Table S18), no significant relationships between task performance and hippocampal grey matter volume were evident.

## 4. DISCUSSION

Marked differences exist across healthy individuals in their ability to imagine visual scenes, recall autobiographical memories, think about the future and navigate in the world. One possible source of this variance may be an individual’s hippocampal grey matter volume. Here, however, in a large sample of young, healthy adult participants, we found little evidence that hippocampal grey matter volume was related to task performance. This was the case across different methodologies (VBM and partial correlations), whole brain or hippocampal ROIs, different sub-groups of participants (divided by gender, task performance and self-reported ability) and when examining latent variables derived from across the cognitive tasks. Hippocampal grey matter volume may not, therefore, significantly influence performance on tasks known to require the hippocampus in healthy people.

Of the 52 main VBM analyses we performed, only two significant associations between hippocampal grey matter volume and task performance were noted. First, a negative association between a cluster of ten voxels and the measure of AI internal place was observed. However, this result was only significant when correcting for the posterior hippocampal mask, and was not replicated when a partial correlation analysis was performed on the extracted hippocampal volume data. Indeed, the correlation coefficient between the posterior hippocampal volume and AI internal place was only −0.053. In the second result, a cluster of seventeen voxels was positively associated with navigation route knowledge when correcting for the anterior hippocampal mask only. However, as shown in Figure 1d, these voxels are clearly on the edge of a larger cluster that was in fact centred on the anterior fusiform and parahippocampal cortices. Overall, given the number of individual analyses performed and the nature of the findings, neither of these results is particularly convincing.

Of the 712 auxiliary analyses performed, only two were significant – AI vividness ratings and the PSIQ by whole hippocampal volume interaction, and AI vividness ratings and the PSIQ by posterior hippocampal volume interaction. Further examination of these interactions found that in individuals with low PSIQ scores (less than 23.65 and 24.13 respectively), there was a negative relationship between whole and posterior hippocampal volume and AI vividness. However, it should be borne in mind that the effect size of the interactions was small (r values of 0.24 and 0.22) and observing only two significant results across the large number of analyses performed is well within the number of false positive results expected, even with statistical correction applied.

The overwhelming number of our analyses produced null results. We are, of course, mindful that null results can be difficult to interpret and that an absence of evidence is not evidence of absence. Nevertheless, several factors motivate confidence in interpreting the current results as showing little evidence of a relationship between hippocampal grey matter volume and performance on our tasks of interest.

First, we had a sample of 217 participants. Samples of this size are much better powered to find true and reliable relationships between brain structure and cognitive ability than the smaller sample sizes (typically between 20 and 30 participants) often tested in previous studies (Kharabian Masouleh et al., 2019; Kong & Francks, 2019). Indeed, the only other study to employ a substantial sample was that of Weisberg et al. (2019; n = 90) who also found no relationship between hippocampal grey matter volume and navigation task performance.

Second, our investigation was wide-ranging and thorough with 26 different outcome measures, each relating to different aspects of scene imagination, autobiographical memory, future thinking and navigation. It seems unlikely that if a relationship existed between hippocampal grey matter volume and these cognitive functions that it would not have been identified by at least one of these performance measures.

Third, our sample was specifically recruited to provide a wide variance in our measures of interest. With the exception of the navigation movie clip recognition sub-measure (where performance was at, or very close to, ceiling) there were large individual differences in ability on all of the tasks. In addition, hippocampal grey matter volume was heterogeneous across the sample. We were, therefore, well-placed to identify any potential relationships between hippocampal volume and task performance, yet few were found.

Fourth, we investigated possible relationships across the whole brain, but also using anatomically-defined hippocampal ROIs. Despite the reduced level of statistical correction required when constraining analyses to hippocampal anatomical ROIs, we still found no significant relationships. It could be argued that the already liberal statistical thresholds should be reduced even further, but this would simply increase the chances of false positive results, given our large sample size and the considerable number of analyses that were performed.

Fifth, similar outcomes were observed using two different analysis methods. While our primary analyses involved VBM, by extracting the whole, anterior and posterior hippocampal volumes for each participant and additionally calculating the posterior:anterior volume ratio, we could also perform partial correlations between each of the volume measurements and our measures of task performance. The results, therefore, do not seem to be consequent upon a specific analysis technique, but instead there was a consistent pattern of null results.

Sixth, we also divided the sample into multiple sub-groups, investigating male and female participants, assessing whether there was any effect of cognitive performance (using both a median split, and by examining only the best and worst performers for each task measure), and self-reported ability. No significant associations between hippocampal volume and performance were identified.

Finally, we also conducted multivariate analyses, using PCA to identify latent variables from across the cognitive tasks. This approach offered increased statistical power and enabled data driven investigations, eschewing a reliance on conceptually driven cognitive constructs, and so had the potential to increase the generalisability of findings beyond the specific tasks in question. To the best of our knowledge, formal PCA analyses using the sub-measures of the tasks employed here have not been reported previously. It was reassuring to find broad correspondence between the components identified by the PCA and the cognitive tasks, and to note that the spatial coherence and the emotional aspects of mental representations cut across cognitive tasks. Of particular relevance for the questions under consideration here, and as with the other analyses, no significant associations between hippocampal grey matter volume and the identified latent variables were observed.

How do our null results correspond to the existing literature? We are not alone in reporting a lack of association between hippocampal grey matter volume and cognitive ability in healthy adults. No previous studies have investigated the hippocampal volume-scene imagination relationship. Considering memory, a meta-analysis of 33 studies identified no relationship between performance on laboratory-based memory tasks and hippocampal volume (Van Petten, 2004). The one previous study that investigated future thinking found no evidence of an association between hippocampal volume and future thinking performance – with voxels instead being located outside of the hippocampus (Yang et al., 2020). Regarding navigation, two previous studies have found no relationship between hippocampal volume and navigational ability in samples drawn from the general population (Maguire, Spiers, et al., 2003; Weisberg et al., 2019). Given the well-known “file drawer” problem (Franco, Malhotra, & Simonovits, 2014; Rosenthal, 1979), the number of null results is likely to be higher than those published in the literature. Our findings, therefore, of no relationship between hippocampal volume and task performance, are not unique to the current sample.

We appreciate that other studies have reported hippocampal volume-task performance relationships. For example, a positive association was identified between posterior hippocampal grey matter volume and recollective memory ability (Poppenk & Moscovitch, 2011), and both posterior and anterior hippocampal grey matter volume have been associated with various measures of spatial navigation, sometimes within specific subgroups (e.g. Bohbot et al., 2007; Brown et al., 2014; Brunec et al., 2019; Chrastil et al., 2017; Hartley & Harlow, 2012; He & Brown, 2020; Schinazi et al., 2013; Sherrill et al., 2018). These disparate results, however, may have been influenced by methodological factors. As alluded to, the sample sizes used were typically small, with the largest being only a quarter of the current study (see Kharabian Masouleh et al., 2019 for more on this issue). Moreover, previous studies have not always been optimally analysed. For example, in some cases relevant covariates were not included, or appropriate levels of statistical correction were not applied, or results were found only in specific smaller sub-groups. Any of these issues could have affected the reliability of the observed results.

The current results differ from those involving licensed London taxi drivers, where increased posterior, but decreased anterior, hippocampal grey matter volume has been consistently identified compared to a variety of control groups, including the same individuals tested before and after they acquired The Knowledge (Maguire et al., 2000; Maguire, Woollett, et al., 2006; Woollett et al., 2008; Woollett & Maguire, 2011). The training to become a London taxi driver takes 3-4 years and approximately 50% of trainees fail to qualify. Those who successfully qualify as London taxi drivers are, therefore, a unique population with extreme spatial navigation expertise. Consequently, the divergence between our null results and those in London taxi drivers is not surprising. Indeed, interpreting the current null results (and those of Maguire, Spiers, et al., 2003, and Weisberg et al. 2019) together with the London taxi driver findings may help to refine our understanding of the relationship between hippocampal volume and navigation ability. Together these results suggest that measurable hippocampal volume changes associated with ability may only be apparent following extreme spatial training rather than being a marker of ability in the general population.

Why might measureable hippocampal volume changes only be detected in the context of extreme expertise? As we did not address this question in the current study we can only speculate. Taking the example of navigation, the vast majority of humans never need to acquire the huge amount of spatial layout knowledge associated with being a licensed London taxi driver. Hence, the hippocampus has likely evolved to accommodate representations of environments that we use day-to-day, with sufficient capacity for a wide range of navigation experiences. Moreover, proficiency, especially at the extreme end of the scale, may come at a cost. For instance, in licensed London taxi drivers, grey matter volume is increased in the posterior hippocampus relative to control groups, but this is accompanied by decreased anterior hippocampal volume (Maguire et al., 2000; Maguire, Woollett, & Spiers, 2006), and also poorer performance on some anterograde visuospatial memory testes (Woollett & Maguire, 2009; 2012; Woollett, Spiers, & Maguire, 2009). Consequently, hippocampal volume may reflect a balance between sufficient functionality for the activities that a typical human undertakes whilst not compromising other hippocampal operations in the service of extreme expertise.

While we found that hippocampal volume seemed to be unrelated to task performance in the general population, it is likely that some neural variations exist that can, at least in part, explain the marked differences in performance observed in tasks such as those we employed. One possibility is that while volumetric differences for the whole hippocampus, or even its anterior or posterior portions, are not related to performance, effects might be observed for the volume of specific hippocampal subfields. For example, larger CA3 volume has been associated with reduced subjective confusion when recalling highly similar memories (Chadwick, Bonnici, & Maguire, 2014). In addition, subiculum volume and combined left dentate gyrus/CA2,3 volume has been positively associated with the number of internal details on the autobiographical interview (Palombo et al., 2018). However, while these studies highlight the potential for relationships between hippocampal subfield volumes and task performance, they also suffer from the limitation of small sample sizes, and so additional work is required to address this issue further.

Another possibility is that the volume of brain regions other than the hippocampus are related to performance on our tasks of interest. Regions including, but not limited to, the parahippocampal, retrosplenial, posterior cingulate, parietal, and medial prefrontal cortices have been associated with scene imagination, autobiographical memory, future thinking and navigation (e.g. Ciaramelli et al., 2017; Hassabis, Kumaran, & Maguire, 2007; Ramanan et al., 2018; Schacter et al., 2012; Simons, Peers, Mazuz, Berryhill, & Olson, 2010; Stawarczyk & D’Argembeau, 2015). However, as reported in the Supplementary Results, when using a p < 0.05 FWE statistical threshold across the whole brain we observed only one relationship – a negative association between 22 voxels in the right parahippocampal cortex and AI vividness ratings. More in-depth analyses using specific ROIs may highlight other relationships (e.g. the positive association between the parahippocampal/fusiform cortices and navigation route knowledge sub-measure we observed when using an uncorrected threshold), but these ROIs would need to be motivated in advance to justify their use, going beyond the remit of the current study.

Advances in neuroimaging mean it is now possible to investigate brain structure in more detail than just grey matter volume. Quantitative neuroimaging techniques (Weiskopf, Mohammadi, Lutti, & Callaghan, 2015) can be used to model different properties of tissue microstructure, in particular that of myelination and iron. Studies using these techniques have, for example, found relationships between myelination and iron in ageing (Callaghan et al., 2014) and, in older participants, increased myelination and decreased iron tissue content has been associated with verbal memory performance in the ventral striatum (Steiger, Weiskopf, & Bunzeck, 2016). Tissue microstructure, both at the whole hippocampus level or within specific subfields, or even in regions outside of the hippocampus, may offer explanations about the variance observed in scene imagination, autobiographical memory, future thinking and navigation ability and will be an interesting target for future work.

The connectivity between brain regions may also be an important factor in explaining individual differences in task performance. Structural measures of connectivity, for example diffusion-weighted imaging of hippocampal white matter pathways, have been associated with both autobiographical memory and navigation ability (Hodgetts et al., 2017; Hodgetts et al., 2020; Iaria, Lanyon, Fox, Giaschi, & Barton, 2008; Metzler-Baddeley, Jones, Belaroussi, Aggleton, & O’Sullivan, 2011). Moreover, functional connectivity between the medial temporal lobe and parietal-occipital regions may be related to questionnaire measures of autobiographical memory (Sheldon, Farb, Palombo, & Levine, 2016), while the functional connectivity of the posterior hippocampus and retrosplenial cortex has been linked with responses on navigation questionnaires (Sulpizio, Boccia, Guariglia, & Galati, 2016). Both structural and functional connectivity measures may, therefore, be associated with task performance, perhaps to a greater extent than hippocampal grey matter volume, and represent important avenues to explore in future studies.

As with functional connectivity, patterns of activity during rest or task-based fMRI may be related to individual differences in performance. For example, higher vividness ratings when performing mental imagery tasks have been reported to correlate with activity in the fusiform, posterior cingulate, parahippocampal (Fulford et al., 2018) and visual (Dijkstra, Bosch, & van Gerven, 2017; Keogh, Bergmann, & Pearson, 2020) cortices. In relation to autobiographical memory, more vivid recall has been linked with higher levels of hippocampal (Gilboa, Winocur, Grady, Hevenor, & Moscovitch, 2004) and precuneus (Richter, Cooper, Bays, & Simons, 2016) activity, while superior memory precision has been related to increased angular gyrus engagement during task performance (Richter et al., 2016). Better performance during navigation tasks has also been associated with increased hippocampal activity (Hartley, Maguire, Spiers, & Burgess, 2003; Ohnishi, Matsuda, Hirakata, & Ugawa, 2006). Individual differences in ability may, therefore, be detectable in fMRI activity during task performance although, as with the structural MRI studies outlined in Section 1, there is a need for large-scale functional neuroimaging studies to examine this further.

## 5. CONCLUSION

In this study we attempted to circumvent issues known to affect studies of healthy people that assess relationships between hippocampal grey matter volume and performance on cognitive tasks that require the hippocampus. Having tested a large sample of healthy participants with a wide range of ability, no credible associations were identified. Consequently, variability in hippocampal grey matter volume seems unlikely to explain individual differences in scene imagination, autobiographical memory, future thinking and spatial navigation performance in the general population. It may be that in healthy participants it is only in extreme situations, as in the case of licensed London taxi drivers, that measurable ability-related hippocampus volume changes are exhibited.

## DECLARTION OF COMPETING INTEREST

The authors declare no competing interests.

## DATA AVAILABILITY

Data and materials will be open-access once the construction of a dedicated data-sharing portal has been finalised. In the meantime, requests for data and materials can be sent to.

## CRediT AUTHORSHIP CONTRIBUTION STATEMENT

**Ian A. Clark:** Conceptualization, Methodology, Investigation, Formal analysis, Writing - original draft, Writing - review & editing. **Anna M. Monk:** Investigation, Writing - review & editing. **Victoria Hotchin:** Investigation, Writing - review & editing. **Gloria Pizzamiglio:** Investigation, Writing - review & editing. **Alice Liefgreen:** Investigation, Writing - review & editing. **Martina F. Callaghan:** Formal analysis, Writing - review & editing. **Eleanor A. Maguire:** Conceptualization, Methodology, Formal analysis, Writing - original draft, Writing - review & editing

## Clark et al. Supplementary Materials

### Supplementary Methods

**Table S1.** Double scoring of the scene construction task.

|  | Rating |  |  |  |  |
| --- | --- | --- | --- | --- | --- |
|  | Spatial<br>References | Entities<br>Present | Sensory<br>Descriptions | Thoughts/<br>Emotions/Actions | Quality<br>Ratings |
| **For each individual scene** |  |  |  |  |  |
| n = 308 | 0.90 | 0.96 | 0.94 | 0.90 | 0.90 |
| **For each individual participant (i.e. score is averaged across the seven scenes)** |  |  |  |  |  |
| n = 44 | 0.91 | 0.99 | 0.97 | 0.91 | 0.93 |

*Note.* Inter-class correlation coefficients from a two way random effect model looking for absolute agreement for each content score and for the quality ratings. Four experimenters scored the whole data set (n = 217 participants, 1519 individual scenes) with double scoring performed on 20% of the data (n = 44 participants, 308 scenes) proportionally for each original experimenter.

**Table S2.** Double scoring of the autobiographical interview.

|  | Rating |  |  |  |  |  |  |
| --- | --- | --- | --- | --- | --- | --- | --- |
|  | Internal<br>Event | Internal<br>Place | Internal<br>Time | Internal<br>Perceptual | Internal<br>Emotion | Internal<br>Sum | External<br>Sum |
| **For each individual memory** |  |  |  |  |  |  |  |
| n = 215 | 0.92 | 0.85 | 0.94 | 0.92 | 0.86 | 0.94 | 0.84 |
| **For each individual participant (i.e. score is averaged across the five memories)** |  |  |  |  |  |  |  |
| n = 43 | 0.95 | 0.88 | 0.96 | 0.94 | 0.81 | 0.97 | 0.87 |

*Note.* Inter-class correlation coefficients from a two way random effects model looking for absolute agreement for each score on the autobiographical interview. Three experimenters scored the whole data set (n = 217 participants, 1085 individual memories) and double scoring was performed 20% of the data (n = 43 participants, 215 individual memories) proportionally for each original experimenter.

**Table S3.** Double scoring of the future thinking task.

|  | Rating |  |  |  |  |
| --- | --- | --- | --- | --- | --- |
|  | Spatial<br>References | Entities<br>Present | Sensory<br>Descriptions | Thoughts/<br>Emotions/Actions | Quality<br>Ratings |
| **For each individual scene** |  |  |  |  |  |
| n = 132 | 0.90 | 0.94 | 0.93 | 0.88 | 0.90 |
| **For each individual participant (i.e. score is averaged across the three future scenes)** |  |  |  |  |  |
| n = 44 | 0.94 | 0.95 | 0.96 | 0.88 | 0.92 |

*Note.* Inter-class correlation coefficients from a two way random effects model looking for absolute agreement for each content score and for the quality ratings. Four experimenters scored the whole data set (n = 217 participants, 651 individual future scenes) with double scoring performed on 20% of the data (n = 44 participants, 132 future scenes) proportionally for each original experimenter.

**Table S4.**
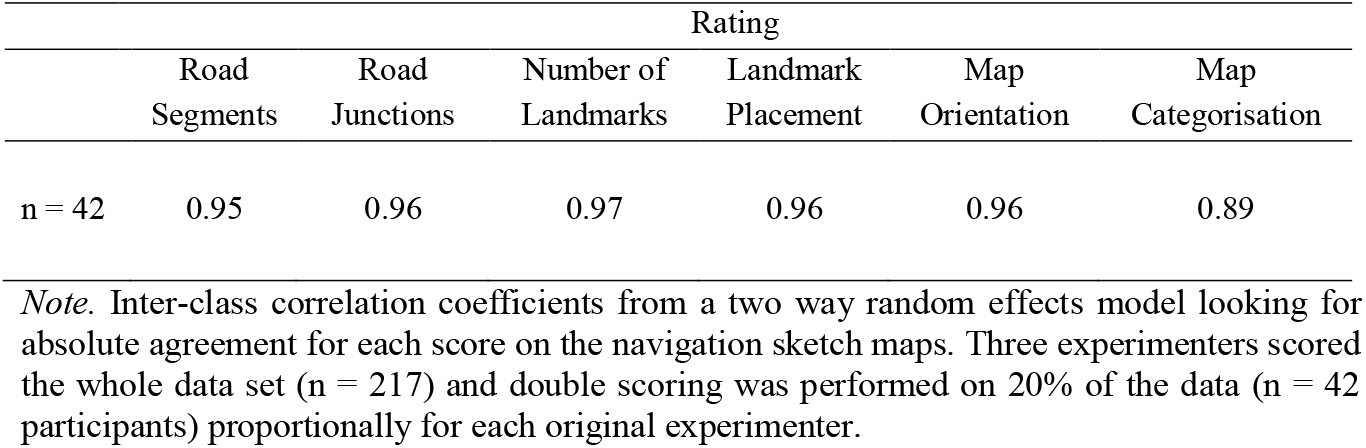
Double scoring of the navigation sketch maps.

|  | Rating |  |  |  |  |  |
| --- | --- | --- | --- | --- | --- | --- |
|  | Road<br>Segments | Road<br>Junctions | Number of<br>Landmarks | Landmark<br>Placement | Map<br>Orientation | Map<br>Categorisation |
| n = 42 | 0.95 | 0.96 | 0.97 | 0.96 | 0.96 | 0.89 |

*Note.* Inter-class correlation coefficients from a two way random effects model looking for absolute agreement for each score on the navigation sketch maps. Three experimenters scored the whole data set (n = 217) and double scoring was performed on 20% of the data (n = 42 participants) proportionally for each original experimenter.

### Supplementary Results

#### Primary VBM analyses: Results outside of the hippocampus

Only one association was identified between cognitive task performance and grey matter volume outside of the hippocampus (when using a statistical threshold of p < 0.05 FWE whole brain corrected). A negative association was observed between 22 voxels in the right parahippocampal cortex and AI vividness ratings (peak coordinates = 24 −23 −28, peak t = 4.78, p _FWE whole brain corrected_ = 0.025).

**Table S5.** Partial correlations between task performance and hippocampal grey matter volume with age, gender, total intracranial volume and MRI scanner as covariates.

| Performance variable | Whole hippocampus |  | Anterior hippocampus |  | Posterior hippocampus |  | Posterior/Anterior hippocampus ratio |  |
| --- | --- | --- | --- | --- | --- | --- | --- | --- |
|  | r | p | r | p | r | p | r | p |
| **Scene construction** |  |  |  |  |  |  |  |  |
| Experiential index | 0.054 | 0.89 | 0.061 | 0.89 | 0.037 | 0.89 | -0.021 | 0.89 |
| Spatial references | 0.082 | 0.89 | 0.079 | 0.89 | 0.068 | 0.89 | 0.00 | 1.00 |
| Entities present | -0.013 | 0.89 | -0.062 | 0.89 | 0.034 | 0.89 | 0.10 | 0.89 |
| Sensory descriptions | 0.023 | 0.89 | 0.069 | 0.89 | -0.024 | 0.89 | -0.095 | 0.89 |
| Thoughts/emotions/actions | -0.043 | 0.89 | -0.066 | 0.89 | -0.013 | 0.89 | 0.037 | 0.89 |
| Spatial coherence index | 0.029 | 0.89 | 0.033 | 0.89 | 0.018 | 0.89 | -0.021 | 0.89 |
| **Autobiographical interview** |  |  |  |  |  |  |  |  |
| Internal details | -0.054 | 0.87 | -0.004 | 0.95 | -0.089 | 0.71 | -0.11 | 0.69 |
| External details | -0.013 | 0.92 | -0.067 | 0.87 | 0.039 | 0.87 | 0.10 | 0.69 |
| Internal events | -0.069 | 0.87 | -0.014 | 0.92 | -0.11 | 0.69 | -0.11 | 0.69 |
| Internal time | -0.027 | 0.87 | -0.025 | 0.87 | -0.024 | 0.87 | -0.013 | 0.92 |
| Internal place | -0.054 | 0.87 | -0.044 | 0.87 | -0.053 | 0.87 | -0.031 | 0.87 |
| Internal perceptual | -0.005 | 0.95 | 0.036 | 0.87 | -0.042 | 0.87 | -0.091 | 0.71 |
| Internal thoughts/emotions | -0.033 | 0.87 | -0.027 | 0.87 | -0.032 | 0.87 | -0.027 | 0.87 |
| Vividness rating | -0.13 | 0.69 | -0.095 | 0.71 | -0.14 | 0.69 | -0.072 | 0.87 |
| **Future thinking** |  |  |  |  |  |  |  |  |
| Experiential index | 0.029 | 0.89 | 0.064 | 0.89 | -0.010 | 0.89 | -0.075 | 0.89 |
| Spatial references | 0.068 | 0.89 | 0.11 | 0.89 | 0.020 | 0.89 | -0.071 | 0.89 |
| Entities present | -0.021 | 0.89 | -0.017 | 0.89 | -0.021 | 0.89 | -0.010 | 0.89 |
| Sensory descriptions | -0.016 | 0.89 | 0.026 | 0.89 | -0.051 | 0.89 | -0.083 | 0.89 |
| Thoughts/emotions/actions | -0.034 | 0.89 | -0.051 | 0.89 | -0.011 | 0.89 | 0.025 | 0.89 |
| Spatial coherence index | -0.025 | 0.89 | -0.011 | 0.89 | -0.031 | 0.89 | -0.037 | 0.89 |
| **Navigation** |  |  |  |  |  |  |  |  |
| Overall navigation score | 0.071 | 0.55 | 0.12 | 0.36 | 0.015 | 0.91 | -0.10 | 0.36 |
| Movie clip recognition | 0.11 | 0.36 | 0.057 | 0.62 | 0.14 | 0.36 | 0.10 | 0.36 |
| Scene recognition | -0.18 | 0.91 | -0.016 | 0.91 | -0.016 | 0.91 | -0.002 | 0.98 |
| Proximity judgements | 0.082 | 0.46 | 0.060 | 0.61 | 0.084 | 0.46 | 0.030 | 0.88 |
| Route knowledge | 0.11 | 0.36 | 0.16 | 0.36 | 0.042 | 0.78 | -0.11 | 0.36 |
| Sketch map | 0.060 | 0.61 | 0.11 | 0.36 | 0.004 | 0.98 | -0.10 | 0.36 |

Note. P values are Benjamini-Hochberg false discovery rate corrected at $p < 0.05$ .

**Table S6.** Partial correlations between task performance and hippocampal grey matter volume in the <u>male</u> participants with age, total intracranial volume and MRI scanner as covariates.

| Performance variable | Whole hippocampus |  | Anterior hippocampus |  | Posterior hippocampus |  | Posterior/Anterior hippocampus ratio |  |
| --- | --- | --- | --- | --- | --- | --- | --- | --- |
|  | r | p | r | p | r | p | r | p |
| **Scene construction** |  |  |  |  |  |  |  |  |
| Experiential index | -0.054 | 0.83 | -0.073 | 0.83 | -0.023 | 0.89 | 0.056 | 0.83 |
| Spatial references | -0.08 | 0.83 | -0.083 | 0.83 | -0.059 | 0.83 | 0.025 | 0.89 |
| Entities present | -0.11 | 0.78 | -0.17 | 0.59 | -0.032 | 0.89 | 0.14 | 0.72 |
| Sensory descriptions | -0.12 | 0.78 | -0.049 | 0.83 | 0.16 | 0.59 | -0.13 | 0.72 |
| Thoughts/emotions/actions | -0.1 | 0.83 | -0.17 | 0.59 | -0.006 | 0.95 | 0.17 | 0.59 |
| Spatial coherence index | 0.034 | 0.89 | 0.006 | 0.95 | 0.053 | 0.83 | 0.058 | 0.83 |
| **Autobiographical interview** |  |  |  |  |  |  |  |  |
| Internal details | -0.086 | 0.79 | -0.039 | 0.84 | -0.11 | 0.79 | -0.10 | 0.79 |
| External details | -0.037 | 0.84 | -0.080 | 0.79 | 0.015 | 0.91 | 0.090 | 0.79 |
| Internal events | -0.038 | 0.84 | 0.022 | 0.90 | -0.088 | 0.79 | -0.12 | 0.79 |
| Internal time | -0.12 | 0.79 | -0.12 | 0.79 | -0.096 | 0.79 | 0.010 | 0.92 |
| Internal place | -0.081 | 0.79 | -0.055 | 0.82 | -0.089 | 0.79 | -0.071 | 0.82 |
| Internal perceptual | -0.11 | 0.79 | -0.053 | 0.82 | -0.14 | 0.79 | -0.12 | 0.79 |
| Internal thoughts/emotions | -0.024 | 0.90 | -0.063 | 0.82 | 0.02 | 0.90 | 0.068 | 0.82 |
| Vividness rating | -0.079 | 0.79 | -0.10 | 0.79 | -0.036 | 0.84 | 0.060 | 0.82 |
| **Future thinking** |  |  |  |  |  |  |  |  |
| Experiential index | -0.12 | 0.52 | -0.056 | 0.68 | -0.15 | 0.52 | -0.11 | 0.52 |
| Spatial references | -0.12 | 0.52 | -0.013 | 0.90 | -0.19 | 0.52 | -0.20 | 0.52 |
| Entities present | -0.11 | 0.52 | -0.094 | 0.55 | -0.11 | 0.52 | -0.024 | 0.87 |
| Sensory descriptions | -0.092 | 0.55 | -0.041 | 0.78 | -0.12 | 0.52 | -0.087 | 0.55 |
| Thoughts/emotions/actions | -0.12 | 0.52 | -0.14 | 0.52 | -0.066 | 0.67 | 0.062 | 0.67 |
| Spatial coherence index | -0.11 | 0.52 | -0.11 | 0.52 | -0.085 | 0.55 | 0.021 | 0.87 |
| **Navigation** |  |  |  |  |  |  |  |  |
| Overall navigation score | 0.07 | 0.77 | 0.13 | 0.46 | -0.005 | 0.97 | -0.14 | 0.43 |
| Movie clip recognition | 0.24 | 0.09 | 0.16 | 0.38 | 0.27 | 0.072 | 0.15 | 0.41 |
| Scene recognition | 0.12 | 0.48 | 0.084 | 0.67 | 0.12 | 0.48 | 0.061 | 0.81 |
| Proximity judgements | 0.012 | 0.97 | 0.017 | 0.97 | 0.004 | 0.97 | -0.011 | 0.97 |
| Route knowledge | 0.18 | 0.35 | 0.27 | 0.072 | 0.044 | 0.88 | -0.24 | 0.090 |
| Sketch map | 0.044 | 0.88 | 0.10 | 0.55 | -0.025 | 0.97 | -0.14 | 0.43 |

Note. P values are Benjamini-Hochberg false discovery rate corrected at $p < 0.05$ .

**Table S7.** Partial correlations between tasks performance and hippocampal grey matter volume in the <u>female</u> participants with age, total intracranial volume and MRI scanner as covariates.

| Performance variable | Whole hippocampus |  | Anterior hippocampus |  | Posterior hippocampus |  | Posterior/Anterior hippocampus ratio |  |
| --- | --- | --- | --- | --- | --- | --- | --- | --- |
|  | r | p | r | p | r | p | r | p |
| **Scene construction** |  |  |  |  |  |  |  |  |
| Experiential index | 0.16 | 0.38 | 0.21 | 0.22 | 0.083 | 0.77 | -0.1 | 0.77 |
| Spatial references | 0.21 | 0.22 | 0.23 | 0.22 | 0.16 | 0.38 | -0.027 | 0.87 |
| Entities present | 0.058 | 0.77 | 0.028 | 0.87 | 0.072 | 0.77 | 0.064 | 0.77 |
| Sensory descriptions | 0.16 | 0.38 | 0.19 | 0.28 | 0.096 | 0.77 | -0.066 | 0.77 |
| Thoughts/emotions/actions | -0.03 | 0.87 | 0.007 | 0.94 | -0.055 | 0.77 | -0.076 | 0.77 |
| Spatial coherence index | 0.014 | 0.93 | 0.056 | 0.77 | -0.026 | 0.87 | -0.097 | 0.77 |
| **Autobiographical interview** |  |  |  |  |  |  |  |  |
| Internal details | -0.035 | 0.98 | 0.029 | 0.98 | -0.082 | 0.91 | -0.13 | 0.89 |
| External details | 0.015 | 0.98 | -0.052 | 0.94 | 0.069 | 0.94 | 0.12 | 0.89 |
| Internal events | -0.098 | 0.91 | -0.049 | 0.94 | -0.12 | 0.89 | -0.10 | 0.91 |
| Internal time | 0.049 | 0.94 | 0.062 | 0.94 | 0.029 | 0.98 | -0.034 | 0.98 |
| Internal place | -0.003 | 0.98 | -0.002 | 0.98 | -0.004 | 0.98 | -0.004 | 0.98 |
| Internal perceptual | 0.066 | 0.94 | 0.11 | 0.89 | 0.013 | 0.98 | -0.086 | 0.91 |
| Internal thoughts/emotions | -0.074 | 0.94 | -0.006 | 0.98 | -0.12 | 0.89 | -0.13 | 0.89 |
| Vividness rating | -0.18 | 0.89 | -0.086 | 0.91 | -0.22 | 0.89 | -0.18 | 0.89 |
| **Future thinking** |  |  |  |  |  |  |  |  |
| Experiential index | 0.23 | 0.12 | 0.24 | 0.12 | 0.17 | 0.34 | -0.03 | 0.88 |
| Spatial references | 0.25 | 0.12 | 0.23 | 0.12 | 0.22 | 0.12 | 0.051 | 0.88 |
| Entities present | 0.12 | 0.64 | 0.11 | 0.64 | 0.10 | 0.64 | 0.008 | 0.95 |
| Sensory descriptions | 0.081 | 0.72 | 0.12 | 0.64 | 0.033 | 0.88 | -0.079 | 0.72 |
| Thoughts/emotions/actions | 0.036 | 0.88 | 0.037 | 0.88 | 0.028 | 0.88 | -0.006 | 0.95 |
| Spatial coherence index | 0.062 | 0.85 | 0.11 | 0.64 | 0.012 | 0.95 | -0.098 | 0.64 |
| **Navigation** |  |  |  |  |  |  |  |  |
| Overall navigation score | 0.044 | 0.97 | 0.079 | 0.97 | 0.005 | 0.97 | -0.066 | 0.97 |
| Movie clip recognition | -0.009 | 0.97 | -0.045 | 0.97 | 0.024 | 0.97 | 0.058 | 0.97 |
| Scene recognition | -0.12 | 0.97 | -0.10 | 0.97 | -0.12 | 0.97 | -0.043 | 0.97 |
| Proximity judgements | 0.16 | 0.97 | 0.12 | 0.97 | 0.16 | 0.97 | 0.059 | 0.97 |
| Route knowledge | 0.015 | 0.97 | 0.015 | 0.97 | 0.012 | 0.97 | 0.009 | 0.97 |
| Sketch map | 0.050 | 0.97 | 0.091 | 0.97 | 0.004 | 0.97 | -0.079 | 0.97 |

Note. P values are Benjamini-Hochberg false discovery rate corrected at $p < 0.05$ .

**Table S8.**
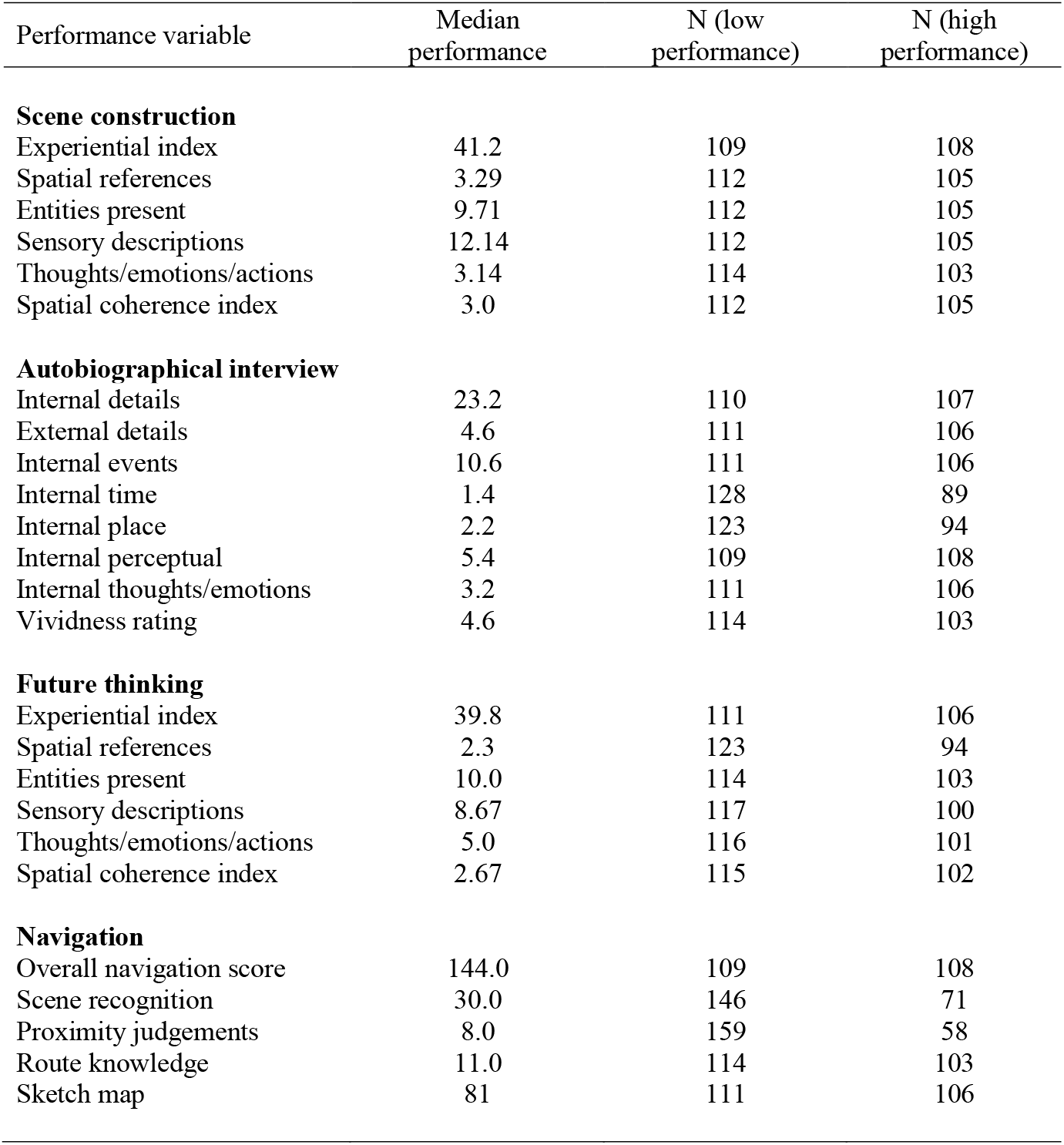
Details of the groups created for each task when dividing by median performance.

| Performance variable | Median performance | N (low performance) | N (high performance) |
| --- | --- | --- | --- |
| **Scene construction** |  |  |  |
| Experiential index | 41.2 | 109 | 108 |
| Spatial references | 3.29 | 112 | 105 |
| Entities present | 9.71 | 112 | 105 |
| Sensory descriptions | 12.14 | 112 | 105 |
| Thoughts/emotions/actions | 3.14 | 114 | 103 |
| Spatial coherence index | 3.0 | 112 | 105 |
| **Autobiographical interview** |  |  |  |
| Internal details | 23.2 | 110 | 107 |
| External details | 4.6 | 111 | 106 |
| Internal events | 10.6 | 111 | 106 |
| Internal time | 1.4 | 128 | 89 |
| Internal place | 2.2 | 123 | 94 |
| Internal perceptual | 5.4 | 109 | 108 |
| Internal thoughts/emotions | 3.2 | 111 | 106 |
| Vividness rating | 4.6 | 114 | 103 |
| **Future thinking** |  |  |  |
| Experiential index | 39.8 | 111 | 106 |
| Spatial references | 2.3 | 123 | 94 |
| Entities present | 10.0 | 114 | 103 |
| Sensory descriptions | 8.67 | 117 | 100 |
| Thoughts/emotions/actions | 5.0 | 116 | 101 |
| Spatial coherence index | 2.67 | 115 | 102 |
| **Navigation** |  |  |  |
| Overall navigation score | 144.0 | 109 | 108 |
| Scene recognition | 30.0 | 146 | 71 |
| Proximity judgements | 8.0 | 159 | 58 |
| Route knowledge | 11.0 | 114 | 103 |
| Sketch map | 81 | 111 | 106 |

**Table S9.** Comparison of hippocampal grey matter volumes when dividing the sample into two groups determined by their median performance on each cognitive task, with age, gender, total intracranial volume and MRI scanner included as covariates.

| Performance variable | Whole hippocampus |  | Anterior hippocampus |  | Posterior hippocampus |  | Posterior/Anterior hippocampus ratio |  |
| --- | --- | --- | --- | --- | --- | --- | --- | --- |
|  | F | p | F | p | F | p | F | p |
| **Scene construction** |  |  |  |  |  |  |  |  |
| Experiential index | 0.85 | 0.70 | 0.48 | 0.83 | 0.88 | 0.70 | 0.051 | 0.99 |
| Spatial references | 1.09 | 0.70 | 0.95 | 0.70 | 0.79 | 0.70 | 0.002 | 0.99 |
| Entities present | 0.043 | 0.99 | 1.06 | 0.70 | 0.35 | 0.83 | 2.93 | 0.70 |
| Sensory descriptions | 0.88 | 0.70 | 0.013 | 0.99 | 2.27 | 0.70 | 2.45 | 0.70 |
| Thoughts/emotions/actions | 1.30 | 0.70 | 1.26 | 0.70 | 0.85 | 0.70 | 0.020 | 0.99 |
| Spatial coherence index | 0.031 | 0.99 | 0.11 | 0.99 | 0.00 | 0.99 | 0.39 | 0.83 |
| **Autobiographical interview** |  |  |  |  |  |  |  |  |
| Internal details | 0.96 | 0.81 | 0.007 | 0.95 | 2.59 | 0.70 | 4.18 | 0.62 |
| External details | 0.53 | 0.81 | 3.65 | 0.62 | 0.26 | 0.81 | 5.81 | 0.54 |
| Internal events | 0.76 | 0.81 | 0.005 | 0.95 | 2.06 | 0.78 | 2.69 | 0.70 |
| Internal time | 0.27 | 0.81 | 0.29 | 0.81 | 0.16 | 0.88 | 0.004 | 0.95 |
| Internal place | 0.13 | 0.89 | 0.009 | 0.95 | 0.50 | 0.81 | 1.17 | 0.81 |
| Internal perceptual | 0.071 | 0.94 | 0.042 | 0.95 | 0.42 | 0.81 | 1.44 | 0.81 |
| Internal thoughts/emotions | 0.63 | 0.81 | 0.27 | 0.81 | 0.77 | 0.81 | 0.58 | 0.81 |
| Vividness ratings | 1.56 | 0.81 | 0.69 | 0.81 | 1.90 | 0.78 | 0.57 | 0.81 |
| **Future thinking** |  |  |  |  |  |  |  |  |
| Experiential index | 0.11 | 0.96 | 0.17 | 0.96 | 0.033 | 0.96 | 0.15 | 0.96 |
| Spatial references | 1.50 | 0.96 | 2.71 | 0.96 | 0.34 | 0.96 | 0.58 | 0.96 |
| Entities present | 0.009 | 0.96 | 0.055 | 0.96 | 0.003 | 0.96 | 0.13 | 0.96 |
| Sensory descriptions | 0.062 | 0.96 | 1.58 | 0.96 | 0.54 | 0.96 | 3.97 | 0.96 |
| Thoughts/emotions/actions | 0.17 | 0.96 | 0.11 | 0.96 | 1.03 | 0.96 | 1.87 | 0.96 |
| Spatial coherence index | 0.005 | 0.96 | 0.006 | 0.96 | 0.039 | 0.96 | 0.020 | 0.96 |
| **Navigation** |  |  |  |  |  |  |  |  |
| Overall navigation score | 0.11 | 0.96 | 0.49 | 0.96 | 0.008 | 0.96 | 0.74 | 0.96 |
| Scene recognition | 0.098 | 0.96 | 0.23 | 0.96 | 0.009 | 0.96 | 0.19 | 0.96 |
| Proximity judgements | 0.079 | 0.96 | 0.046 | 0.96 | 0.081 | 0.96 | 0.003 | 0.96 |
| Route knowledge | 0.72 | 0.96 | 1.76 | 0.96 | 0.052 | 0.96 | 1.12 | 0.96 |
| Sketch map | 0.11 | 0.96 | 0.29 | 0.96 | 0.007 | 0.96 | 0.21 | 0.96 |

Note. P values are Benjamini-Hochberg false discovery rate corrected at $p < 0.05$ .

**Table S10.** Partial correlations between task performance and hippocampal grey matter volume in the <u>low</u> performing participants only (as determined by a median split for each task) with age, gender, total intracranial volume and MRI scanner as covariates.

| Performance variable | Whole hippocampus |  | Anterior hippocampus |  | Posterior hippocampus |  | Posterior/Anterior hippocampus ratio |  |
| --- | --- | --- | --- | --- | --- | --- | --- | --- |
|  | r | p | r | p | r | p | r | p |
| **Scene construction** |  |  |  |  |  |  |  |  |
| Experiential index | 0.002 | 0.98 | 0.059 | 0.98 | -0.054 | 0.98 | 0.10 | 0.96 |
| Spatial references | 0.019 | 0.98 | -0.003 | 0.98 | 0.036 | 0.98 | 0.052 | 0.98 |
| Entities present | 0.013 | 0.98 | -0.005 | 0.98 | 0.027 | 0.98 | 0.042 | 0.98 |
| Sensory descriptions | 0.17 | 0.88 | 0.18 | 0.88 | 0.14 | 0.96 | -0.008 | 0.98 |
| Thoughts/emotions/actions | -0.057 | 0.98 | -0.11 | 0.96 | 0.008 | 0.98 | 0.11 | 0.96 |
| Spatial coherence index | 0.053 | 0.98 | 0.097 | 0.96 | -0.007 | 0.98 | -0.10 | 0.96 |
| **Autobiographical interview** |  |  |  |  |  |  |  |  |
| Internal details | -0.032 | 0.91 | -0.022 | 0.96 | -0.033 | 0.91 | -0.004 | 0.97 |
| External details | -0.082 | 0.90 | -0.047 | 0.90 | -0.10 | 0.90 | -0.065 | 0.90 |
| Internal events | -0.072 | 0.90 | -0.020 | 0.96 | -0.11 | 0.90 | -0.11 | 0.90 |
| Internal time | 0.047 | 0.90 | 0.006 | 0.97 | 0.073 | 0.90 | 0.064 | 0.90 |
| Internal place | -0.049 | 0.90 | -0.075 | 0.90 | -0.014 | 0.97 | 0.042 | 0.90 |
| Internal perceptual | -0.054 | 0.90 | -0.099 | 0.90 | 0.010 | 0.97 | 0.13 | 0.90 |
| Internal thoughts/emotions | 0.047 | 0.90 | -0.034 | 0.91 | 0.12 | 0.90 | 0.16 | 0.90 |
| Vividness ratings | -0.11 | 0.90 | -0.077 | 0.90 | -0.11 | 0.90 | -0.067 | 0.90 |
| **Future thinking** |  |  |  |  |  |  |  |  |
| Experiential index | 0.075 | 0.99 | 0.14 | 0.99 | 0.001 | 0.99 | -0.11 | 0.99 |
| Spatial references | 0.032 | 0.99 | 0.008 | 0.99 | 0.047 | 0.99 | 0.063 | 0.99 |
| Entities present | -0.003 | 0.99 | 0.019 | 0.99 | -0.023 | 0.99 | -0.039 | 0.99 |
| Sensory descriptions | 0.056 | 0.99 | 0.061 | 0.99 | 0.041 | 0.99 | -0.010 | 0.99 |
| Thoughts/emotions/actions | -0.069 | 0.99 | -0.008 | 0.99 | -0.11 | 0.99 | -0.14 | 0.99 |
| Spatial coherence index | -0.049 | 0.99 | 0.040 | 0.99 | -0.12 | 0.99 | -0.19 | 0.99 |
| **Navigation** |  |  |  |  |  |  |  |  |
| Overall navigation score | 0.29 | 0.06 | 0.24 | 0.065 | 0.27 | 0.06 | 0.067 | 0.60 |
| Scene recognition | -0.077 | 0.55 | -0.094 | 0.49 | -0.044 | 0.67 | 0.055 | 0.60 |
| Proximity judgements | 0.082 | 0.52 | 0.054 | 0.60 | 0.094 | 0.48 | 0.057 | 0.60 |
| Route knowledge | 0.18 | 0.19 | 0.16 | 0.24 | 0.15 | 0.24 | 0.032 | 0.78 |
| Sketch map | 0.25 | 0.065 | 0.22 | 0.084 | 0.21 | 0.090 | 0.020 | 0.84 |

Note. P values are Benjamini-Hochberg false discovery rate corrected at $p < 0.05$ .

**Table S11.** Partial correlations between performance and hippocampal grey matter volume in the <u>high</u> performing participants only (as determined by a median split for each task) with age, gender, total intracranial volume and MRI scanner as covariates.

| Performance variable | Whole hippocampus |  | Anterior hippocampus |  | Posterior hippocampus |  | Posterior/Anterior hippocampus ratio |  |
| --- | --- | --- | --- | --- | --- | --- | --- | --- |
|  | r | p | r | p | r | p | r | p |
| **Scene construction** |  |  |  |  |  |  |  |  |
| Experiential index | -0.019 | 0.99 | -0.001 | 0.99 | -0.032 | 0.99 | -0.033 | 0.99 |
| Spatial references | 0.057 | 0.99 | 0.067 | 0.99 | 0.036 | 0.99 | -0.027 | 0.99 |
| Entities present | -0.017 | 0.99 | -0.021 | 0.99 | -0.009 | 0.99 | 0.012 | 0.99 |
| Sensory descriptions | 0.078 | 0.99 | 0.066 | 0.99 | 0.071 | 0.99 | -0.001 | 0.99 |
| Thoughts/emotions/actions | 0.096 | 0.99 | 0.062 | 0.99 | 0.11 | 0.99 | 0.072 | 0.99 |
| Spatial coherence index | 0.014 | 0.99 | -0.072 | 0.99 | 0.080 | 0.99 | 0.18 | 0.99 |
| **Autobiographical interview** |  |  |  |  |  |  |  |  |
| Internal details | 0.0 | 1.0 | 0.004 | 1.0 | -0.003 | 1.0 | -0.003 | 1.0 |
| External details | 0.11 | 1.0 | 0.12 | 1.0 | 0.076 | 1.0 | -0.023 | 1.0 |
| Internal events | -0.016 | 1.0 | -0.027 | 1.0 | -0.001 | 1.0 | 0.029 | 1.0 |
| Internal time | -0.067 | 1.0 | 0.004 | 1.0 | -0.12 | 1.0 | -0.16 | 1.0 |
| Internal place | -0.087 | 1.0 | -0.13 | 1.0 | -0.037 | 1.0 | 0.10 | 1.0 |
| Internal perceptual | 0.047 | 1.0 | 0.12 | 1.0 | -0.018 | 1.0 | -0.13 | 1.0 |
| Internal thoughts/emotions | -0.040 | 1.0 | 0.005 | 1.0 | -0.072 | 1.0 | -0.11 | 1.0 |
| Vividness ratings | -0.093 | 1.0 | -0.087 | 1.0 | -0.079 | 1.0 | -0.013 | 1.0 |
| **Future thinking** |  |  |  |  |  |  |  |  |
| Experiential index | -0.027 | 0.91 | 0.017 | 0.91 | -0.061 | 0.91 | -0.082 | 0.91 |
| Spatial references | -0.012 | 0.91 | 0.035 | 0.91 | -0.054 | 0.91 | -0.11 | 0.91 |
| Entities present | -0.012 | 0.91 | -0.064 | 0.91 | 0.040 | 0.91 | 0.10 | 0.91 |
| Sensory descriptions | -0.11 | 0.91 | -0.16 | 0.91 | -0.047 | 0.91 | 0.097 | 0.91 |
| Thoughts/emotions/actions | -0.099 | 0.91 | -0.089 | 0.91 | -0.088 | 0.91 | -0.022 | 0.91 |
| Spatial coherence index | -0.066 | 0.91 | -0.073 | 0.91 | -0.046 | 0.91 | 0.023 | 0.91 |
| **Navigation** |  |  |  |  |  |  |  |  |
| Overall navigation score | -0.071 | 0.69 | 0.039 | 0.74 | -0.15 | 0.33 | -0.22 | 0.24 |
| Scene recognition | 0.11 | 0.63 | 0.16 | 0.40 | 0.038 | 0.76 | -0.14 | 0.47 |
| Proximity judgements | 0.24 | 0.33 | 0.21 | 0.33 | 0.20 | 0.33 | 0.06 | 0.73 |
| Route knowledge | 0.044 | 0.73 | 0.15 | 0.33 | -0.06 | 0.73 | -0.21 | 0.26 |
| Sketch map | -0.082 | 0.63 | 0.048 | 0.73 | -0.18 | 0.33 | -0.27 | 0.12 |

Note. P values are Benjamini-Hochberg false discovery rate corrected at $p < 0.05$ .

**Table S12.** Details of the groups created for each task when taking only the best and worst performers.

| Performance variable | Worst performers maximum | Best performers minimum | N (worst performers) | N (best performers) |
| --- | --- | --- | --- | --- |
| **Scene construction** |  |  |  |  |
| Experiential index | 32.66 | 48.49 | 20 | 20 |
| Spatial references | 1.57 | 5.57 | 21 | 22 |
| Entities present | 6.29 | 13.86 | 20 | 20 |
| Sensory descriptions | 7.71 | 17.0 | 21 | 21 |
| Thoughts/emotions/actions | 1.29 | 6.14 | 19 | 18 |
| Spatial coherence index | 0.71 | 5.0 | 23 | 25 |
| **Autobiographical interview** |  |  |  |  |
| Internal details | 15.20 | 34.60 | 22 | 21 |
| External details | 1.80 | 10.20 | 21 | 21 |
| Internal events | 6.20 | 16.60 | 21 | 21 |
| Internal time | 2.20 | 10.0 | 18 | 15 |
| Internal place | 1.40 | 3.0 | 26 | 32 |
| Internal perceptual | 0.40 | 2.60 | 20 | 21 |
| Internal thoughts/emotions | 1.20 | 6.0 | 19 | 18 |
| Vividness rating | 3.40 | 5.60 | 19 | 20 |
| **Future thinking** |  |  |  |  |
| Experiential index | 30.13 | 47.33 | 21 | 21 |
| Spatial references | 0.67 | 4.33 | 32 | 30 |
| Entities present | 6.67 | 14.67 | 23 | 22 |
| Sensory descriptions | 4.33 | 13.33 | 21 | 21 |
| Thoughts/emotions/actions | 2.33 | 8.33 | 22 | 22 |
| Spatial coherence index | 0.0 | 5.0 | 23 | 29 |
| **Navigation** |  |  |  |  |
| Overall navigation score | 95.0 | 191.0 | 21 | 21 |
| Scene recognition | 27.0 | 32.0 | 34 | 28 |
| Proximity judgements | 5.0 | 10.0 | 18 | 13 |
| Route knowledge | 6.0 | 19.0 | 23 | 22 |
| Sketch map | 39.0 | 121.0 | 21 | 21 |

**Table S13.** Comparison of hippocampal grey matter volumes when taking the <u>best and worst</u> performing participants for task, with age, gender, total intracranial volume and MRI scanner included as covariates.

| Performance variable | Whole hippocampus |  | Anterior hippocampus |  | Posterior hippocampus |  | Posterior/Anterior hippocampus ratio |  |
| --- | --- | --- | --- | --- | --- | --- | --- | --- |
|  | F | p | F | p | F | p | F | p |
| **Scene construction** |  |  |  |  |  |  |  |  |
| Experiential index | 0.092 | 0.97 | 0.36 | 0.97 | 0.0 | 1.0 | 0.39 | 0.97 |
| Spatial references | 0.69 | 0.97 | 0.15 | 0.97 | 1.24 | 0.97 | 1.34 | 0.97 |
| Entities present | 0.0 | 1.0 | 0.38 | 0.97 | 0.37 | 0.97 | 1.94 | 0.97 |
| Sensory descriptions | 0.47 | 0.97 | 0.63 | 0.97 | 0.27 | 0.97 | 0.006 | 1.0 |
| Thoughts/emotions/actions | 0.014 | 1.0 | 0.21 | 0.97 | 0.49 | 0.97 | 1.29 | 0.97 |
| Spatial coherence index | 0.22 | 0.97 | 0.34 | 0.97 | 0.075 | 0.97 | 0.058 | 0.97 |
| **Autobiographical interview** |  |  |  |  |  |  |  |  |
| Internal details | 4.09 | 0.80 | 1.17 | 0.88 | 6.04 | 0.61 | 2.44 | 0.83 |
| External details | 0.73 | 0.88 | 1.36 | 0.88 | 0.089 | 0.88 | 0.69 | 0.88 |
| Internal events | 1.56 | 0.88 | 0.30 | 0.88 | 2.62 | 0.83 | 1.67 | 0.88 |
| Internal time | 0.001 | 0.98 | 0.053 | 0.90 | 0.035 | 0.91 | 0.23 | 0.88 |
| Internal place | 0.48 | 0.88 | 0.10 | 0.88 | 0.85 | 0.88 | 0.58 | 0.88 |
| Internal perceptual | 0.18 | 0.88 | 0.13 | 0.88 | 0.17 | 0.88 | 0.092 | 0.88 |
| Internal thoughts/emotions | 0.51 | 0.88 | 0.87 | 0.88 | 0.17 | 0.88 | 0.43 | 0.88 |
| Vividness ratings | 2.89 | 0.98 | 1.85 | 0.88 | 2.42 | 0.83 | 0.097 | 0.88 |
| **Future thinking** |  |  |  |  |  |  |  |  |
| Experiential index | 0.17 | 0.96 | 0.80 | 0.96 | 0.001 | 0.98 | 0.85 | 0.96 |
| Spatial references | 0.74 | 0.96 | 3.06 | 0.96 | 0.001 | 0.98 | 1.75 | 0.96 |
| Entities present | 0.20 | 0.96 | 0.029 | 0.98 | 0.37 | 0.96 | 0.44 | 0.96 |
| Sensory descriptions | 0.003 | 0.98 | 0.074 | 0.96 | 0.12 | 0.96 | 0.43 | 0.96 |
| Thoughts/emotions/actions | 0.30 | 0.96 | 0.44 | 0.96 | 0.12 | 0.96 | 0.062 | 0.96 |
| Spatial coherence index | 0.72 | 0.96 | 0.24 | 0.96 | 1.02 | 0.96 | 0.50 | 0.96 |
| **Navigation** |  |  |  |  |  |  |  |  |
| Overall navigation score | 0.88 | 0.47 | 4.51 | 0.09 | 0.071 | 0.79 | 8.39 | 0.06 |
| Scene recognition | 0.58 | 0.56 | 0.93 | 0.47 | 0.21 | 0.72 | 0.25 | 0.72 |
| Proximity judgements | 5.38 | 0.07 | 2.57 | 0.22 | 6.19 | 0.067 | 1.84 | 0.32 |
| Route knowledge | 7.34 | 0.07 | 6.78 | 0.065 | 4.30 | 0.09 | 6.15 | 0.067 |
| Sketch map | 0.99 | 0.47 | 5.18 | 0.073 | 0.098 | 0.79 | 10.01 | 0.06 |

Note. P values are Benjamini-Hochberg false discovery rate corrected at $p < 0.05$ .

**Table S14.** Partial correlations between task performance and the interaction between hippocampal grey matter volume and self-reported ability (measured by the Plymouth Sensory Imagery Questionnaire for scene construction, autobiographical memory and future thinking and the Santa Barbara Sense of Direction Scale for navigation), with age, gender, total intracranial volume and MRI scanner included as covariates in all analyses, as well as the additional covariates of the relevant questionnaire and hippocampal volume measurement.

| Performance variable | Whole hippocampus |  | Anterior hippocampus |  | Posterior hippocampus |  | Posterior/Anterior hippocampus ratio |  |
| --- | --- | --- | --- | --- | --- | --- | --- | --- |
|  | r | p | r | p | r | p | r | p |
| **Scene construction** |  |  |  |  |  |  |  |  |
| Experiential index | 0.044 | 0.94 | 0.054 | 0.94 | 0.029 | 0.94 | -0.023 | 0.94 |
| Spatial references | -0.029 | 0.94 | -0.01 | 0.94 | -0.047 | 0.94 | -0.022 | 0.94 |
| Entities present | 0.002 | 0.97 | 0.009 | 0.94 | -0.017 | 0.94 | -0.038 | 0.94 |
| Sensory descriptions | -0.076 | 0.94 | -0.057 | 0.94 | -0.078 | 0.94 | -0.008 | 0.94 |
| Thoughts/emotions/actions | 0.055 | 0.94 | 0.052 | 0.94 | 0.047 | 0.94 | -0.021 | 0.94 |
| Spatial coherence index | 0.015 | 0.94 | 0.019 | 0.94 | 0.01 | 0.94 | -0.021 | 0.94 |
| **Autobiographical interview** |  |  |  |  |  |  |  |  |
| Internal details | 0.014 | 0.87 | 0.018 | 0.87 | 0.017 | 0.87 | -0.022 | 0.87 |
| External details | 0.084 | 0.78 | 0.11 | 0.50 | 0.033 | 0.87 | -0.12 | 0.49 |
| Internal events | 0.029 | 0.87 | 0.033 | 0.87 | 0.031 | 0.87 | -0.032 | 0.87 |
| Internal time | -0.052 | 0.87 | -0.043 | 0.87 | -0.054 | 0.87 | 0.001 | 0.99 |
| Internal place | 0.029 | 0.87 | 0.075 | 0.87 | -0.027 | 0.87 | -0.14 | 0.29 |
| Internal perceptual | -0.034 | 0.87 | -0.015 | 0.87 | -0.042 | 0.87 | -0.028 | 0.87 |
| Internal thoughts/emotions | 0.067 | 0.87 | 0.02 | 0.87 | 0.11 | 0.50 | 0.086 | 0.78 |
| Vividness ratings | 0.22 | 0.016* | 0.18 | 0.096 | 0.24 | 0.013* | 0.026 | 0.87 |
| **Future thinking** |  |  |  |  |  |  |  |  |
| Experiential index | -0.018 | 0.90 | 0.012 | 0.91 | -0.041 | 0.82 | -0.041 | 0.82 |
| Spatial references | -0.07 | 0.82 | -0.028 | 0.82 | -0.1 | 0.82 | -0.051 | 0.82 |
| Entities present | -0.062 | 0.82 | -0.05 | 0.82 | -0.068 | 0.82 | -0.007 | 0.91 |
| Sensory descriptions | -0.12 | 0.82 | -0.09 | 0.82 | -0.13 | 0.82 | -0.033 | 0.82 |
| Thoughts/emotions/actions | 0.066 | 0.82 | 0.076 | 0.82 | 0.042 | 0.82 | -0.072 | 0.82 |
| Spatial coherence index | 0.03 | 0.82 | 0.045 | 0.82 | 0.011 | 0.91 | -0.03 | 0.82 |
| **Navigation** |  |  |  |  |  |  |  |  |
| Overall navigation score | 0.049 | 0.85 | 0.026 | 0.85 | 0.067 | 0.72 | 0.035 | 0.85 |
| Clip recognition | 0.084 | 0.72 | 0.084 | 0.72 | 0.071 | 0.72 | -0.04 | 0.85 |
| Scene recognition | -0.11 | 0.72 | -0.088 | 0.72 | -0.12 | 0.72 | -0.02 | 0.88 |
| Proximity judgements | 0.1 | 0.72 | 0.074 | 0.72 | 0.11 | 0.72 | 0.047 | 0.85 |
| Route knowledge | 0.002 | 0.98 | -0.012 | 0.90 | 0.016 | 0.89 | 0.028 | 0.85 |
| Sketch map | 0.056 | 0.84 | 0.031 | 0.85 | 0.075 | 0.72 | 0.037 | 0.85 |

Note. P values are Benjamini-Hochberg false discovery rate corrected at $p < 0.05$ . \* = $p < 0.05$ .

**Table S15.** Details of the principal component analysis on the four main outcome measures.

| Cognitive task | Component weighting |
| --- | --- |
| Scene construction experiential index | 0.90 |
| Future thinking experiential index | 0.87 |
| Autobiographical interview internal details | 0.66 |
| Navigation | 0.49 |

Note. The factor scores for each participant were extracted using the Anderson-Rubin method. These were included in a VBM analysis with age, gender, scanner and total intracranial volume as covariates, finding no significant relationships with hippocampal volume both at the whole brain level ( $p < 0.05$ FWE corrected) and when using the anatomical hippocampal masks ( $p < 0.05$ FWE corrected for each mask). Performing partial correlations between the extracted hippocampal grey matter volumes and the factor scores also identified no significant relationships (whole hippocampal volume: $r = 0.032$ , $p_{\text{FDR corrected}} = 0.85$ ; anterior hippocampal volume: $r = 0.073$ , $p_{\text{FDR corrected}} = 0.58$ ; posterior hippocampal volume: $r = -0.012$ , $p_{\text{FDR corrected}} = 0.86$ ; posterior/anterior hippocampal volume ratio: $r = -0.091$ , $p_{\text{FDR corrected}} = 0.58$ ).

**Table S16.** Proportion of variance explained by each principal component of the sub-measures principal component analysis.

| PCA Component | Variance explained |
| --- | --- |
| C1. Content scenes and future | 19.11 |
| C2. Navigation | 11.67 |
| C3. Spatial coherence | 9.88 |
| C4. Thoughts/emotions | 9.42 |
| C5. Autobiographical memory | 9.21 |
| C6. Time | 5.73 |

**Table S17.** Details of the principal component analysis conducted on all 21 task sub-measures.

| Sub-measure | Content scenes<br>and future | Navigation | Spatial<br>coherence | Thoughts/<br>emotions | Autobiographical<br>memory | Time |
| --- | --- | --- | --- | --- | --- | --- |
| Future thinking entities present | 0.84 |  |  |  |  |  |
| Future thinking spatial references | 0.78 |  |  |  |  |  |
| Future thinking sensory descriptions | 0.76 |  |  |  |  |  |
| Scene construction sensory descriptions | 0.75 |  |  |  |  |  |
| Scene construction entities present | 0.72 |  |  | 0.41 |  |  |
| Scene construction spatial references | 0.70 |  |  |  |  |  |
| Navigation route knowledge |  | 0.84 |  |  |  |  |
| Navigation sketch map |  | 0.83 |  |  |  |  |
| Navigation movie clip recognition |  | 0.64 |  |  |  |  |
| Navigation scene recognition |  | 0.59 |  |  |  |  |
| Navigation proximity judgements |  | 0.41 |  |  |  |  |
| Scene construction spatial coherence index |  |  | 0.89 |  |  |  |
| Future thinking spatial coherence index |  |  | 0.87 |  |  |  |
| Autobiographical memory vividness |  |  | 0.61 |  |  | 0.38 |
| Scene construction thoughts/emotions/actions |  |  |  | 0.84 |  |  |
| Future thinking thoughts/emotions/actions |  |  |  | 0.76 |  |  |
| Autobiographical memory internal thoughts/emotions |  |  |  | 0.49 | 0.57 |  |
| Autobiographical memory internal events |  |  |  |  | 0.79 |  |
| Autobiographical memory internal perceptual |  |  |  |  | 0.65 |  |
| Autobiographical memory internal place |  |  |  |  | 0.57 | 0.35 |
| Autobiographical memory internal time |  |  |  |  |  | 0.73 |

Note. Task order is for display purposes only. Only values over 0.3 are reported for ease of viewing.

**Table S18.** Partial correlations between the sub-measures principal components and hippocampal grey matter volume with age, gender, total intracranial volume and MRI scanner as covariates.

| PCA component | Whole hippocampus |  | Anterior hippocampus |  | Posterior hippocampus |  | Posterior/Anterior hippocampus ratio |  |
| --- | --- | --- | --- | --- | --- | --- | --- | --- |
|  | r | p | r | p | r | p | r | p |
| C1. Content scenes and future | 0.041 | 0.78 | 0.048 | 0.78 | 0.027 | 0.88 | -0.009 | 0.97 |
| C2. Navigation | 0.12 | 0.65 | 0.13 | 0.65 | 0.085 | 0.66 | -0.029 | 0.88 |
| C3. Spatial coherence | -0.044 | 0.78 | -0.021 | 0.91 | -0.056 | 0.77 | -0.055 | 0.77 |
| C4. Thoughts/emotions | -0.052 | 0.77 | -0.084 | 0.66 | -0.011 | 0.97 | 0.060 | 0.77 |
| C5. Autobiographical memory | -0.052 | 0.77 | 0.002 | 0.98 | -0.091 | 0.66 | -0.12 | 0.65 |
| C6. Time | -0.096 | 0.66 | -0.097 | 0.66 | -0.075 | 0.75 | 0.006 | 0.97 |

Note. P values are Benjamini-Hochberg false discovery rate corrected at $p < 0.05$ .

**Figure S1.**
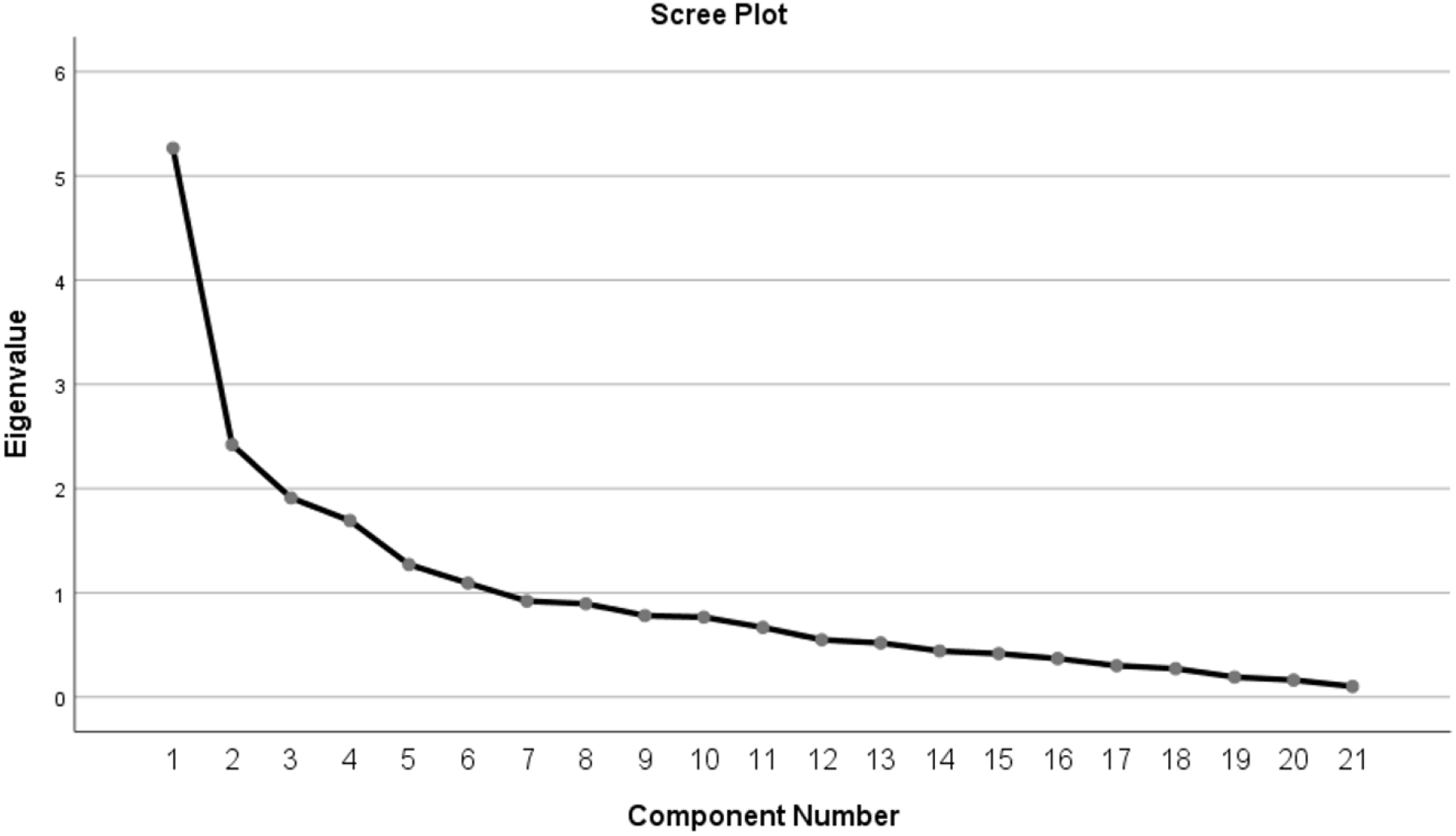
Scree plot of the principal component analysis performed on the sub-measures.

